# Cytosolic Class I J-domain proteins aid mitochondrial protein import and influence homeostasis in *Arabidopsis thaliana*

**DOI:** 10.1101/2024.04.19.590371

**Authors:** Silviya S. Lal, Neha, Yadvendradatta Rajendra Prasad Yadav, Sreehari P., Amit K. Verma, Chandan Sahi

## Abstract

Mitochondrial homeostasis heavily relies on import of numerous nuclear encoded proteins. Multiple chaperone machineries operate at the cytosolic and mitochondrial side of the bilayer to ensure the efficacy of this process. While the mechanisms and components involved in translocation of polypeptides into the mitochondria are known in great detail, the cytosolic events of mitochondrial protein import are poorly understood in plants. This study explores the role of two cytosolic Class I J-domain proteins (JDPs), atDjA1 and atDjA2, in mitochondrial homeostasis in *Arabidopsis thaliana*. We show that atDjA1 and atDjA2 uniquely interacted with outer mitochondrial surface receptors, Tom9, Metaxin, and OM64, which guide the import of numerous mitochondrial precursors. Knockout of atDjA1 or atDjA2 impeded seed germination under multiple abiotic stress conditions, with atDjA1 mutant being more sensitive. Both the mutants demonstrated decreased respiratory rates and enhanced accumulation of reactive oxygen species (ROS), indicating compromised mitochondrial function. Further, knockout of atDjA1 resulted in a significant decline in the number of active mitochondria as well as import of MTS-GFP. While atDjA2 mutants did not show a reduction in mitochondrial number, both the mutants had an activated mitochondrial retrograde signaling (MRS) further underscoring compromised mitochondrial homeostasis in these mutants. Our findings suggest that evolutionarily conserved cytosolic Class I JDPs preserve mitochondrial homeostasis by targeting specific receptors on the outer mitochondrial surface, facilitating the import of cytosolically synthesized mitochondrial precursors in *Arabidopsis thaliana*.

## Introduction

Mitochondrial homeostasis depends on the coordinated activities of proteins encoded by both the nuclear and mitochondrial genomes. Given that nearly 99% of mitochondrial proteins are encoded by the nuclear genome, an efficient import system involving components from both the cytosol and mitochondria is essential (Lister *et al*., 2005; Glaser, Whelan and Logan, 2007; Chacinska *et al*., 2009; Murcha *et al*., 2014). Disruption in mitochondrial function significantly impacts plant’s well-being, leading to substantial impairments in seed germination, growth, flowering, photosynthesis, respiration efficiency, regulation of reactive oxygen species (ROS), and stress sensitivity (Schwarzländer and Finkemeier, 2013; Wang *et al*., 2018; Farooq *et al*., 2021; Van Aken, 2021).

Cytosolic chaperones play crucial roles in the synthesis and maintenance of mitochondrial precursor proteins in an unfolded, import-competent state, preventing their aggregation, thereby ensuring their successful delivery to the mitochondria (Avendaño-Monsalve, Ponce-Rojas and Funes, 2020). Moreover, some cytosolic chaperones facilitate mitochondrial import of precursor proteins by directly interacting with receptors of the outer mitochondrial membrane (Voos, 2003). The Tom70 component of the TOM (Translocase of the Outer Membrane) complex interacts with cytosolic chaperones, including Hsp70 and Hsp90, and carries out mitochondrial protein import (Young, Hoogenraad and Hartl, 2003). Similarly, in *Arabidopsis thaliana*, the cytosolic Hsp90 and Hsp70 bind the outer membrane receptor, OM64, for facilitating mitochondrial protein import (Panigrahi, Whelan and Vrielink, 2014). Membrane receptors are primed for preprotein loading when a chaperone attaches to them. This keeps the receptors open to receiving preproteins and expands the volume of their preprotein-binding pocket (Li *et al*., 2009). After passing through the TOM complex, molecular chaperones in the intermembrane space and mitochondrial matrix take over to facilitate the movement of proteins to different sub-compartments within the mitochondria (Neupert and Pfanner, 1993; Voos and Röttgers, 2002; Adriaenssens *et al*., 2023).

Another class of cytosolic chaperones implicated in mitochondrial import are Hsp40s, also called J-domain proteins (JDPs). They operate as obligatory co-chaperones of Hsp70s, stimulating their intrinsic ATPase activity and facilitating the transfer of diverse protein substrates to Hsp70s, thus driving Hsp70’s multifunctionality (Kampinga and Craig, 2010). Mitochondrial protein import is primarily linked with Class I JDPs, the most abundant type that has all three of the structurally characterized domains identified in the prototypical member, DnaJ of *E. coli*. Their interaction with the Hsp70s is facilitated by a compact tetrahelical J domain, which is connected to four repeats of the CxxCxGxG-type zinc-finger motif (ZnF) through a glycine-rich (G/F) linker region. Following this, there is a carboxy-terminal domain (CTD) consisting of two β-barrel structures, which aids in binding with various client proteins, thus determining their functional specificity (Kampinga and Craig, 2010; Craig and Marszalek, 2017; Kampinga *et al*., 2019). The most abundant Class I JDP in *S. cerevisiae*, Ydj1 facilitates the import of several mitochondrial precursor proteins like Atp2, Porins and F_1_β (Becker *et al*., 1996; Yamamoto *et al*., 2009; Jores *et al*., 2018). Another Class I JDP in budding yeast, Xdj1, has also been implicated in the import of numerous precursors, aiding in mitochondrial biogenesis *via* its interaction with Tom22 (Sahi *et al*., 2013; Opaliński *et al*., 2018). Similarly, mammalian Class I JDPs DnaJA1, 2, and 4 also facilitate the import of proteins to mitochondria (Kanazawa *et al*., 1997; Terada *et al*., 1997; Bhangoo *et al*., 2007). While the translocases found in plant mitochondrial membranes and the process by which pre-protein traverses them are well known (Murcha *et al*., 2014; Ghifari, Gill-Hille and Murcha, 2018), the cytosolic steps and chaperones involved in mitochondrial import are poorly understood in plants.

*A. thaliana* harbors two highly similar cytosolic Class I JDPs, atDjA1 and atDjA2 (alternatively known as AtDjA3/AtJ3/J3 and AtJ2/J2 respectively). Specifically, atDjA1 has been implicated in several housekeeping as well as stress-related functions like abiotic stress tolerance, intracellular pH maintenance and developmental processes such as flowering (Yang *et al*., 2010; Zhou *et al*., 2018; Wang *et al*., 2021). On the other hand, not much is known about atDjA2, except having some unknown, overlapping role with atDjA1 in imparting thermotolerance to Arabidopsis (Li *et al*., 2007). Both atDjA1 and atDjA2 are orthologs of yeast Ydj1 protein (Verma *et al*., 2017). Although possible, the involvement of these cytosolic JDPs in mitochondrial protein import and, therefore, in maintaining mitochondrial homeostasis has not been investigated. Here we examined the role of atDjA1 and atDjA2 in maintaining mitochondrial homeostasis in *A. thaliana*. Purified atDjA1 and atDjA2 differentially interacted with the cytosolic domains of *A. thaliana* outer mitochondrial membrane receptors Tom9.2, Metaxin and OM64. Consistent with a role of atDjA1 and atDjA2 in facilitating the import of precursor proteins from the cytosol to the mitochondria, plants deficient in atDjA1 or atDjA2 showed compromised mitochondrial activity with decreased respiratory efficiency and increased ROS levels, ultimately heightening the vulnerability of these mutants to abiotic stresses. Although, atDjA1 mutant exhibited delayed germination under normal conditions and appeared more susceptible to abiotic stresses than atDjA2 mutant, defects in both atDjA1 and atDjA2 mutants resulted in altered expression of many nuclear genes that regulate Mitochondrial Retrograde Signaling (MRS). Furthermore, the absence of atDjA1 led to a substantial reduction in the number of functional mitochondria as well as a defect in mitochondrial protein import, indicating compromised mitochondrial biogenesis and function. Thus, this study emphasizes the significance of the plant cytosolic JDPs in preserving mitochondrial homeostasis via unique interactions with receptors at the protein entry gates on the outer mitochondrial membrane.

## Results

### atDjA1 and atDjA2 interact with Arabidopsis mitochondrial outer membrane receptor proteins

In budding yeast, the cytosolic Class I JDP, Xdj1 is implicated in the import of nuclear-encoded mitochondrial precursor proteins from the cytoplasm (Sahi *et al*., 2013; Opaliński *et al*., 2018). Further, the interaction between the client binding domain (CTD) domain of Xdj1 and the cytosolic domain of the outer mitochondrial membrane protein Tom22 is required for this function (Opaliński *et al*., 2018) **(Fig. 1A)**. Though *xdj1*Δ cells display no evident growth phenotype, when paired with the loss of Pam17, a component of the mitochondrial inner membrane import machinery, exhibit slow growth as well as mitochondrial import errors at lower temperatures (Sahi *et al*., 2013). As Class I JDPs, both atDjA1 and atDjA2 exhibit a domain organization similar to that of Xdj1 **(Fig. 1B)**. Moreover, the C-terminal peptide binding pockets of atDjA1 and atDjA2 are quite similar to Xdj1 (Verma *et al*., 2017) **(Supplemental Fig. S1)**. While both atDjA1 and atDjA2 rescue the slow growth phenotype of Δ*xdj1*Δ*pam17*, atDjA2 was more efficient in importing Mdj1 into yeast mitochondria (Verma *et al*., 2017). Since the C-terminal domains often determine the functional specificity of JDPs, we hypothesized that the greater efficiency of atDjA2 in complementing Xdj1 functions in budding yeast stems from unique residues in the CTD. To test this, we compared the CTD of Xdj1 (aa 235-377) with atDjA1 (aa 212-370) and atDjA2 (aa 213-370). Three residues, S218, K315 and A366 were identified in atDjA2 that were identical in Xdj1 but different in atDjA1. Among these, S218 and K315 were chosen for mutation and altered to their corresponding residues in atDjA1 -S218P and K315N **(Fig. 1, C and D)**. To ascertain the functional relevance of these unique residues, single and double mutations were created in atDjA2. Comparable growth in the spot assay demonstrated that atDjA2 single mutants (*atDjA2_S218P* and *atDjA2_K315N*) complemented the slow-growth phenotype of *xdj1*Δ*pam17*Δ cells at lower temperature in a manner similar to that of atDjA2 (*data not shown*). However, *atDjA2_S218P+K315N* double, mutants were significantly defective in rescuing the growth defects of *xdj1*Δ*pam17*Δ cells **(Fig. 1E)**. The expression of all the constructs was verified by immunoblot analysis using anti-atDjA1 antibody which recognizes both atDjA1 and atDjA2 proteins equally **(Fig. 1F, Supplemental Fig. S2)**. Additionally, *xdj1*Δ*pam17*Δ cells harboring *atDjA2_S218P+K315N* mutant accumulated the cytosolic Mdj1 precursor protein **(Fig. 1G),** suggesting that the C-terminal client binding domain contributes to the functionality of atDjA2 in substituting for Xdj1 in budding yeast.

**Figure 1.**
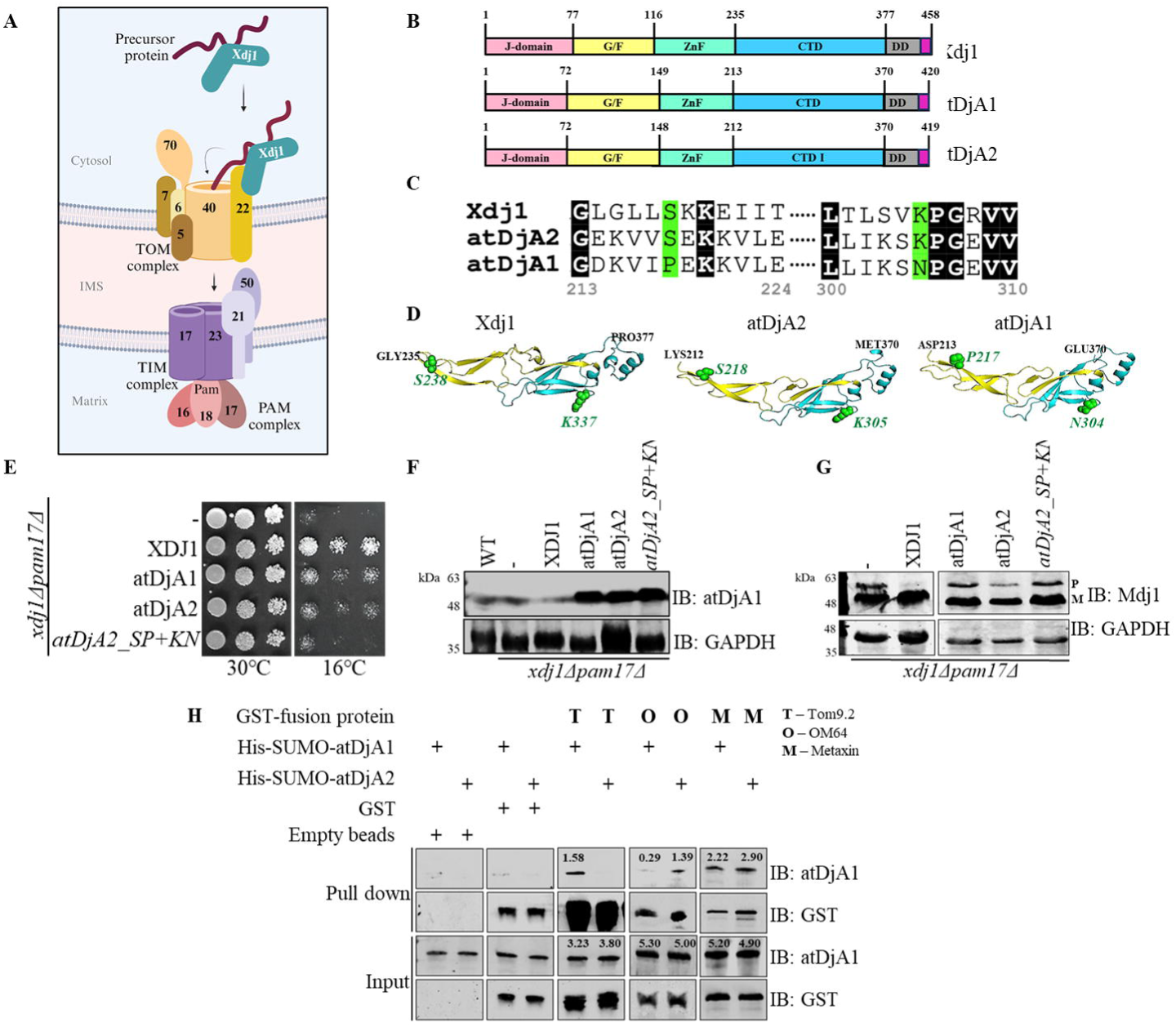
atDjA1 and atDjA2 interact with outer mitochondrial membrane proteins. **A)** Schematic depiction showing the interaction of Xdj1 with outer mitochondrial membrane receptor Tom22 in *S. cerevisiae*, for facilitating mitochondrial import of proteins **B)** Domain organization of *S. cerevisiae* cytosolic Class I JDPs, Ydj1 and Xdj1, and *Arabidopsis thaliana* cytosolic Class I JDPs, atDjA1 and atDjA2. Different domains – J-domain (J), Glycine/Phenylalanine rich region (G/F), Zinc-Finger (ZnF), C-terminal domain (CTD), and Dimerization domain (DD) are shown with its amino acid boundary. The farnesylation motif is shown by pink color at the C-term end. **C)** Amino acid sequence alignment of a portion of the C-terminal domains of Xdj1 (aa 235-342), atDjA2 (aa 213-310) and atDjA1 (aa 212-309) was performed using the MAFFT program. The resulting alignment was subsequently utilized in EsPript to generate the image. The residues that are identical in all three proteins are highlighted in black and the residues in atDjA2 selected for point mutations along with corresponding residues in others are highlighted in green. Starting and ending amino acids are numbered according to atDjA2 sequence position. **D)** Cartoon models of CTD of Xdj1 (aa 235-377), atDjA2 (aa 213-370) and atDjA2 (aa 212-370) created using SWISS-MODEL homology-modelling server using the previously solved structure of Ydj1 (1NLT), followed by further processing in PyMol. Residues selected for mutation in atDjA2 and the corresponding residues in Xdj1 and atDjA1 are shown as green spheres with amino acid labelling. **E)** Equal volumes of *xdj1*Δ*pam17*Δ cells harboring either an empty pRS414-*TEF* vector or JDP expression plasmids [pRS314-*XDJ1*-Xdj1 (XDJ1), pRS414-*TEF*-atDjA1 (atDjA1), pRS414-*TEF*-atDjA2 (atDjA2) together with the atDjA2 mutant, pRS414-*TEF-atDjA2_SP+KN* (*atDjA2_SP+KN*) were spotted on tryptophan drop-out (Trp DO) plates, incubated for three days at 30°C or ten days at 16°C, before being photographed. **F)** Equal amounts of total cell lysate prepared from W303 WT cells and *xdj1*Δ*pam17*Δ cells harboring the above-mentioned JDP expression plasmids were separated by SDS-PAGE, electroblotted onto nitrocellulose membrane and probed with anti-atDjA1 antibody. GAPDH was employed as the loading control. G) Equal amounts of total cell lysate prepared from *xdj1*Δ*pam17*Δ cells harboring either an empty pRS414-*TEF* vector or those harboring the above-mentioned JDP expression plasmids were separated by SDS-PAGE, electroblotted onto nitrocellulose membrane and probed with anti-Mdj1 antibody to identify the cytosolic precursor form (P) and mature mitochondrial form (M) of mitochondrial protein Mdj1. GAPDH was employed as the loading control. **H)** Purified proteins – GST-Tom9.2, GST-OM64, and GST-Metaxin were immobilized on GSH-beads and incubated for 1h at 4°C in GST equilibration buffer with purified His+SUMO-atDjA1/A2. Pre-washing input fractions and post-washing pull-down fractions were separated, resolved by SDS-PAGE, electroblotted onto nitrocellulose membrane and probed with anti-atDjA1 and anti-GST antibodies. The bead-bound GST tag by itself and the empty GSH beads served as negative controls. Band intensities assessed by the ImageJ program are indicated above the bands.

Since both atDjA1 and atDjA2 partially substitute for Xdj1 in yeast, we hypothesized that these might be operating in a similar manner in Arabidopsis, through interactions with outer mitochondrial receptor proteins. So, we explored their physical interaction with known Arabidopsis outer mitochondrial membrane receptor proteins. Three such proteins—Tom9.2, Metaxin, and OM64 were selected, which are known for their roles in facilitating the import of various mitochondrial proteins (Duncan, Murcha and Whelan, 2013). For *in-vitro* interaction studies, the cytosolic domains of Tom9.2, Metaxin, and OM64 proteins, tagged with an N-term GST tag, and full-length atDjA1 and atDjA2 proteins, tagged with an N-terminal His-SUMO tag, were expressed and purified from *E. coli*. Pull-down assays revealed varying affinities of atDjA1 and atDjA2 to different receptor proteins analyzed. While both atDjA1 and atDjA2 were able to associate with the cytosolic domain of Metaxin; only atDjA1 and only atDjA2 could interact with the cytosolic domains of Tom9.2 and OM64, respectively **(Fig. 1H)**. These findings suggest that akin to their yeast counterparts, the ability of atDjA1 and atDjA2 to interact differentially with outer mitochondrial receptor proteins may define their role in mitochondrial protein import and thereby homeostasis in *Arabidopsis thaliana*.

### atDjA1 is required for germination under normal conditions

Next, we set out to decipher the role of atDjA1 and atDjA2 in Arabidopsis. First, the cytosolic localization of both atDjA1 and atDjA2 was confirmed in protoplasts by employing GFP-tagged fusion constructs. In contrast to the poor signals from empty GFP-expressing protoplasts, the GFP fluorescence signals were consistently and robustly visible in the cytosol of GFP-atDjA1/A2 expressing protoplasts **(Supplemental Fig. S3)**. To better comprehend the physiological significance of atDjA1 and atDjA2, we monitored the expression levels of atDjA1 and atDjA2 transcripts in different tissue samples of wild type Arabidopsis, collected at various stages of development, including dry seeds, germinating seeds, seedlings, rosette leaves, cauline leaves, closed buds, flowers, young siliques, senescent siliques, and adult roots. As shown (**Fig. 2, A and B**), transcript encoding atDjA1 and atDjA2 were expressed differentially in every tissue examined, with both being abundantly expressed in siliques and dry seeds, suggesting a conceivable importance of these proteins in the development or germination of seeds. However, on average, atDjA1 was about three times more abundantly expressed than atDjA2 in different tissues **(Fig. 2, A and B)**. The abundance of atDjA1 was further confirmed by assessing the protein levels in endogenous Myc-tagged atDjA1/atDjA2 stable lines **(Supplemental Fig. S4)**.

**Figure 2.**
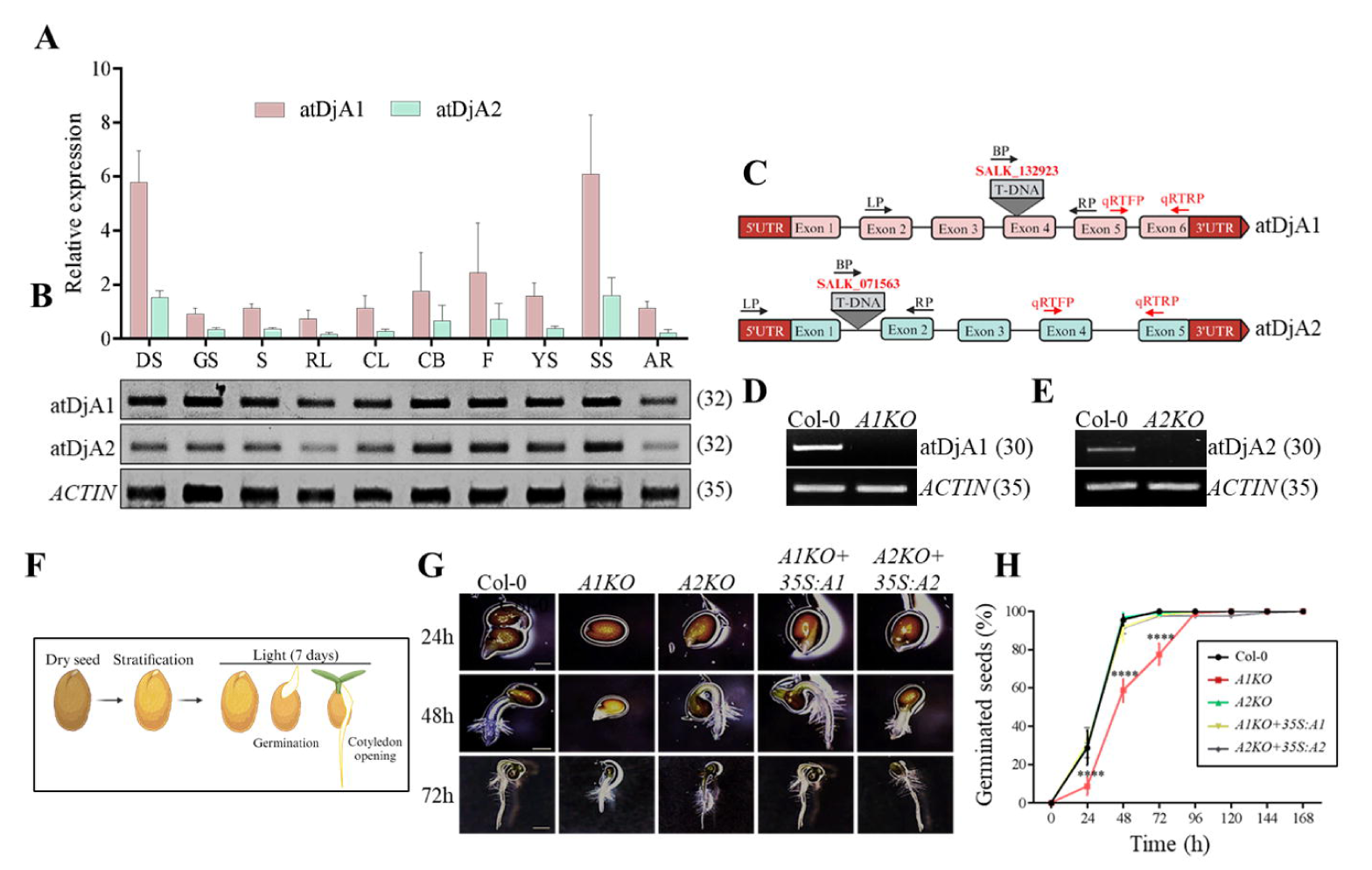
atDjA1 and atDjA2 are differentially expressed. **A)** Expression levels of atDjA1 and atDjA2 genes, in different developmental stages of wild type Col-0 plants grown under long-day conditions including, dry seeds (DS), germinating seeds (GS), 7-day old seedlings (S), rosette leaves (RS), cauline leaves (CL), closed buds (CB), flowers (F), young siliques (YS), senescent siliques (SS) and adult roots (AR), determined by qRT-PCR. The fold change was normalized against Arabidopsis *ACTIN* (AT3G18780) and *EL*α*1* (AT1G07940) genes. The results reflect the mean ± SE of three independent experiments. **B)** Expression levels of atDjA1 and atDjA2 genes, along with *ACTIN* across the indicated tissue samples obtained from Col-0 plants analyzed by semi-quantitative RT-PCR, followed by gel-based analysis. Values in the bracket indicate the number of cycles. **C)** The location of T-DNA inserts and primers for genotyping (black arrows) and qRT-PCR (C-term fragment, red arrows) for atDjA1 and atDjA2 are shown in their respective gene structures. Untranslated region (UTR), T-DNA border primer (BP), left primer (LP), right primer (RP), qRT-forward primer (qRTFP) and qRT-reverse primer (qRTRP). **D-E)** Semiquantitative RT-PCR for **D)** atDjA1 and **E)** atDjA2 in 15-day-old seedlings of WT, *A1KO* and *A2KO* lines (30 cycles). *ACTIN* gene was employed as control (35 cycles). **F)** Schematic depiction of the different phases of seed germination examined in the phenotypic studies. **G)** To compare their rate of germination, seeds of the indicated genotypes were allowed to germinate on ½ MS for 24 hours (24h), 48 hours (48h) and 72 hours (72h) under long-day conditions **H)** Graphs displaying the germination rate of the specified genotypes during a 24-168-hour period on ½ MS plates. The results are based on the mean ± SE of three biological replicates. An asterisk (****, *P<0.0001*, Tukey’s test, One-way ANOVA) indicates a significant difference.

To understand the role of atDjA1 and atDjA2 in Arabidopsis, T-DNA insertion mutant lines for atDjA1: SALK_132923 (insertion in exon 4) and atDjA2: SALK_071563 (insertion in intron 1) were obtained from ABRC **(Fig. 2C)**. These mutants were screened for homozygosity by genotyping PCR **(Supplemental Fig. S5, A and B)**, followed by gene expression analyses by semi-quantitative (full-length transcript) and quantitative (C-terminal fragment) RT-PCR. A substantial reduction in the transcript was detected using C-term specific PCR primers for both atDjA1 (∼70% reduction) and atDjA2 (∼50% reduction) genes in the respective mutant lines as compared to the wild-type (Col-0) **(Supplemental Fig. S5, C, D and E)**. However, the full-length transcript was not detected in either atDjA1 or atDjA2 mutants, suggesting that these lines are knockouts **(Fig. 2, D and E)**. Hereafter, these lines are referred to as *A1KO* (atDjA1 knockout) and *A2KO* (atDjA2 knockout). The *A1KO+35S:A1* and *A2KO+35S:A2* complementation lines were verified for respective transcript levels by qRT-PCR, which were comparable to Col-0 plants **(Supplemental Fig. S5E)**.

Upon confirmation, equally aged seeds of knockout and complementation lines were subjected to comprehensive phenotypic analysis, with emphasis on germination and seedling establishment. The mutant seeds, along with that of Col-0 and respective complemented lines (*A1KO+35S:A1* and *A2KO+35S:A2*) were allowed to germinate on ½ MS with 1% (w/v) sucrose. The plates were kept under long-day conditions, and the number of seedlings displaying radicle emergence and open cotyledons was recorded **(Fig. 2F)**. The assay was performed for a total of seven days, or 168h, at 24h intervals, and the percentage of germination each day was calculated and plotted. Additionally, the percentage of seedlings with open cotyledons on the seventh day was also plotted. On ½ MS medium, the *AIKO* seeds exhibited slower germination till 72h, in comparison with *A2KO* and Col-0 **(Fig. 2G)**. *A1KO* seeds required 48h of light exposure for 50% of their seeds to germinate, while *A2KO* and Col-0 seeds achieved the same in 24h (*P<0.0001*) **(Fig. 2, G and H).** After 72h of light treatment, fewer than 50% of *A1KO* seeds developed green cotyledons, in contrast to 70–80% of seeds from Col-0 and *A2KO* **(Supplemental Fig. S6A)**. This defect in *A1KO* germination was rescued in the *A1KO+35S:A1* complemented line **(Fig. 2, G and H; Supplemental Fig. S6A)**. However, by 72h, nearly all seeds from every genotype had germinated, and on the seventh day of the experiment (7 Days After Germination/ DAG), all of them developed genuine leaves and looked indistinguishable **(Supplemental Fig. S6B)**. These findings imply that atDjA1 plays a key role in germination in *A. thaliana*.

### atDjA1 and atDjA2 are required for germination and seedling establishment under stress conditions

Both atDjA1 and atDjA2 are stress-inducible, with atDjA1 transcript being selectively induced by high temperature and atDjA2 by a variety of abiotic stressors, such as salt and osmotic stresses (Verma *et al*., 2017). To evaluate the function of atDjA1 and atDjA2 under different abiotic stresses, the mutants along with respective complemented lines were allowed to germinate on ½ MS media supplemented with gradually increasing concentrations of NaCl (50 mM, 75 mM, 100 mM and 150 mM), which induce both ionic and osmotic stress, and mannitol (100 mM, 200 mM, 250 mM and 300 mM) and glucose (2%, 4% and 6%), which induce osmotic stress. Higher concentrations of all three stressors adversely affected seed germination as well as the rate of cotyledon development of both *A1KO* and *A2KO* lines **(Fig. 3)**.

**Figure 3.**
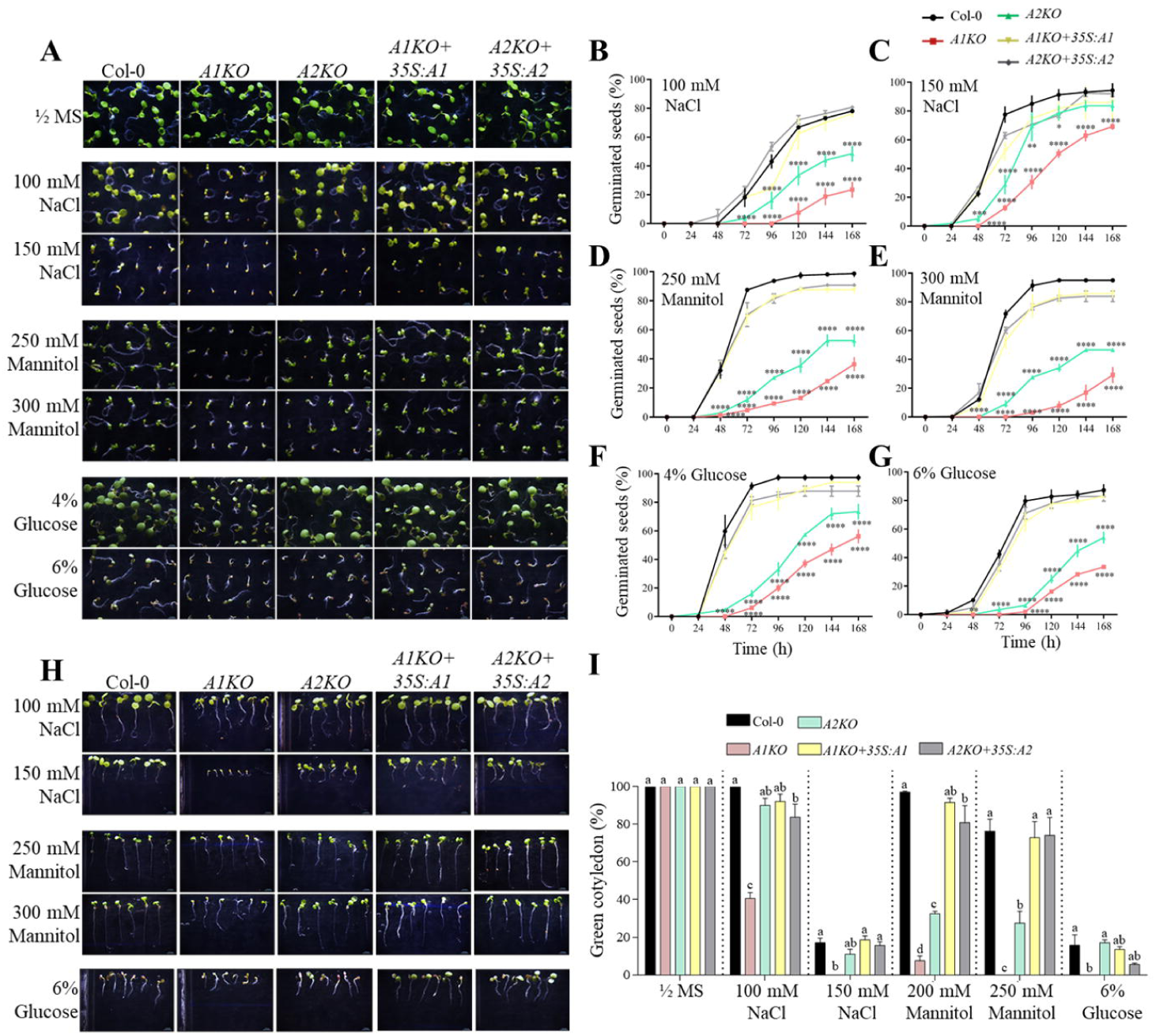
Loss of atDjA1 and atDjA2 impairs seed germination under different stress conditions. **A)** Seeds of the indicated genotypes were allowed to germinate for 7 days on ½ MS medium and that supplemented with indicated concentrations of NaCl (100 mM and 150 mM), mannitol (250 mM and 300 mM) and glucose (4% and 6%). **B-G)** Graphs displaying the germination rate of the specified genotypes during a 24-168-hour period on **B)** 100 mM NaCl, **C)** 150 mM NaCl, **D)** 250 mM mannitol, **E)** 300 mM mannitol, **F)** 4% glucose and **G)** 6% glucose. The results are based on the mean ± SE of three biological replicates. An asterisk (****, *P<0.0001*; ***, *P<0.001*; **, *P<0.01* and *, *P<0.05*, Tukey’s test, One-way ANOVA) indicates a significant difference. **H)** Representative photographs of 7-day-old seedlings of different genotypes under indicated concentrations of NaCl (100 mM and 150 mM), mannitol (250 mM and 300 mM) and glucose (6%). **I)** Graphical representation of the percentage of open green cotyledons for the indicated genotypes counted on the seventh day of analysis. Results are based on the mean ± SE of three biological replicates. Columns with different letters indicate significant differences (Tukey’s test, Two-way ANOVA).

On 50 mM NaCl, *A1KO* seeds required longer (96h) than *A2KO* and Col-0 (48h) to germinate 50% of their seeds **(Supplemental Fig. S7, A and B)**. On 75 mM NaCl, *A1KO* seeds achieved 50% germination only after 120h of light exposure and A*2KO* seeds after 72h, in contrast to 48h for Col-0. The delay in germination rate was rescued effectively by the *A1KO+35S:A1* and *A2KO+35S:A2* complemented lines **(Supplemental Fig. S7, A and C)**. The total germination on the seventh day for both *A1KO* and *A2KO* on 75 mM NaCl was also significantly lower (60-70% for *A1KO* and 80-85% for *A2KO*) **(Supplemental Fig. S7C)**. On 100 mM NaCl, both *A1KO* and *A2KO* seeds germinated more slowly, with only 60–70% of the *AIKO* seeds, 80-85% of the *A2KO* seeds as well as about 95% of the Col-0 and complementation seeds **(Fig. 3, A and B)**. Also, only 40% of seeds developed open cotyledons in *A1KO* lines, compared to nearly 100% in others **(Fig. 3, H and I)**. 150 mM NaCl had a severe effect on both the germination and cotyledon development of both *A1KO* and *A2KO*. A total germination of only 20-25% for *A1KO*, as opposed to 45-50% for *A2KO* and 75-80% for Col-0 and complemented lines were observed **(Fig. 3, A and C)**. *A1KO* additionally demonstrated a severe impairment in post-germination development, with no open cotyledons, as contrasted with 15-20% seeds with open cotyledons in *A2KO*, Col-0 and complemented lines **(Fig. 3, H and I)**.

Both *A1KO* and *A2KO* seeds revealed reduced radicle emergence and cotyledon opening as compared to Col-0 when sown on all concentrations of osmotic stressors tested, from 100 mM to 300 mM mannitol **(Fig. 3; Supplemental Fig. S8)** and 2% to 6% glucose **(Fig. 3; Supplemental Fig. S9)**. On 100 mM mannitol, *A1KO* seeds achieved 50% germination in 120h, versus 72h by *A2KO* seeds, as compared to 48h by Col-0 **(Supplemental Fig. S8, A and B)**. *A1KO* seeds achieved an overall germination of 75– 80% at the end of the experiment, whereas the other genotypes had almost 100% germination **(Supplemental Fig. S8B)**. *A1KO* and *A2KO* seeds exhibited severe defects when subjected to 200 mM mannitol with a total germination of 45–50% for *A1KO* seeds and 75–80% for *A2KO* seeds, in comparison with 95–100% germination in Col-0 and complemented lines **(Supplemental Fig. S8, A and C)**. On plates containing 250 mM and 300 mM mannitol, the germination and seedling establishment of *A1KO* and *A2KO* seedlings were both severely hindered. In comparison to 85–100% of Col-0 and complemented seedlings, only 40% of *A1KO* and 50% of *A2KO* seeds germinated on 250 mM mannitol after seven days **(Fig. 3, A and D)**. Of those, only 5–10% of *A1KO* and 30–35% of *A2KO* seeds had open cotyledons **(Fig. 3, H and I)**. On 300 mM mannitol-containing plates, only 25–30% of *A1KO* and 40–45% of *A2KO* seeds germinated **(Fig. 3, A and E)**. Also, *A1KO* failed to show any true open cotyledons, although *A2KO* did in as little as 30% of seeds, compared to 70–80% of seeds in Col-0 **(Fig. 3, H and I)**.

Another osmotic stressor, glucose, when added in increasing concentrations to ½ MS medium (without sucrose), impacted seed germination, caused post-germinative growth halt, and prevented cotyledon expansion and greening in all genotypes **(Fig. 3; Supplemental Fig. S9)**. On 2% glucose plates, *A1KO* seeds required 120h and *A2KO* seeds required 72h to achieve 50% germination, however, Col-0 and complemented lines just required 48h **(Supplemental Fig. S9)**. For *A1KO*, only a total of 80% seeds germinated at the end of the assay, compared to 95-100% in others, with all the seedlings displaying open green cotyledons **(Supplemental Fig. S9B)**. In the presence of 4% glucose, both *A1KO* and *A2KO* had delayed germination; a total of only 50% of seeds germinated in *A1KO*, 70% in *A2KO* and 95% in Col-0 and complemented lines **(Fig. 3, A and F)**. The effects of 6% glucose on the phenotypes of *A1KO* and *A2KO* seeds were extremely adverse. This included significantly delayed germination, with just 30% of seeds in *A1KO* seeds and 50% in *A2KO* seeds germinating overall, compared to 90–95% in Col-0 and complemented lines **(Fig. 3G)**. Additionally, the cotyledon opening was also hampered, with no open cotyledons for *A1KO* and 15-20% in *A2KO*, Col-0, and complemented lines. The color of the developing cotyledons ranged from pale green to yellow for all genotypes **(Fig. 3, H and I)**. In all the examined conditions, the complemented lines (*A1KO+35S:A1* and *A2KO+35S:A2*) behaved comparable to Col-0, except for a slight delay in germination at higher concentrations of mannitol and glucose. This implies that the reduced levels of atDjA1 and atDjA2 are responsible for both the germination and post-germination defects observed in the presence of these stressors. In all the conditions analyzed, *A1KO* lines often showed higher vulnerability to in terms of both delayed germination and stalled cotyledon development as compared to *A2KO* lines. Together, these findings reveal the significance of the Class I JDPs atDjA1 and atDjA2 for early germination and seedling establishment under various stressful conditions.

### Knockout of atDjA1 and atDjA2 results in reduced respiration

For germination and seedling establishment to be efficient, respiration and the associated generation of energy are necessary (Botha, Potgieter and Botha, 1992). Therefore, impairment in mitochondrial activity impacts the oxygen consumption rate, resulting in anomalies associated with seed germination and post-germination development (Farooq *et al*., 2021). To investigate if the reduced amounts of atDjA1 or atDjA2 affected seed respiration, measurements of oxygen consumption by germinating seeds of *A1KO* and *A2KO* lines were performed using a Clark-type electrode. Both stratified seeds (0h light) and seeds that undergo germination (48h light) were studied for their rates of total oxygen absorption. In 0h light-treated imbibed seeds, the amount of oxygen consumption by the mutants was comparable to that of Col-0 seeds **(Fig. 4A)**. Seeds from all the genotypes exhibited a consistent increase in respiration rate as compared to 0h, when the germination process commenced under the light treatment. However, the amount of oxygen consumed by *A1KO* and *A2KO* seeds was significantly less as compared to the Col-0 seeds. Following 48h of light treatment, both *A1KO* and *A2KO* germinating seeds consumed two times less oxygen than Col-0 (*P<0.0001* for *A1KO* and *P<0.001* for *A2KO*) **(Fig. 4A)**. Additionally, the Oxygen Consumption Graph (OCR) graph plotted over a 10 min time frame suggested a reduced oxygen consumption rate in *A1KO* and *A2KO* germinating seeds than Col-0 **(Fig. 4B)**. As expected, the complemented lines *A1KO+35S:A1* and *A2KO+35S:A2* successfully regained the mutants’ decreased respiration rate, implying that the oxygen consumption defect in mutant seeds is a consequence of lower atDjA1 and atDjA2 levels **(Fig. 4, A and B)**.

**Figure 4.**
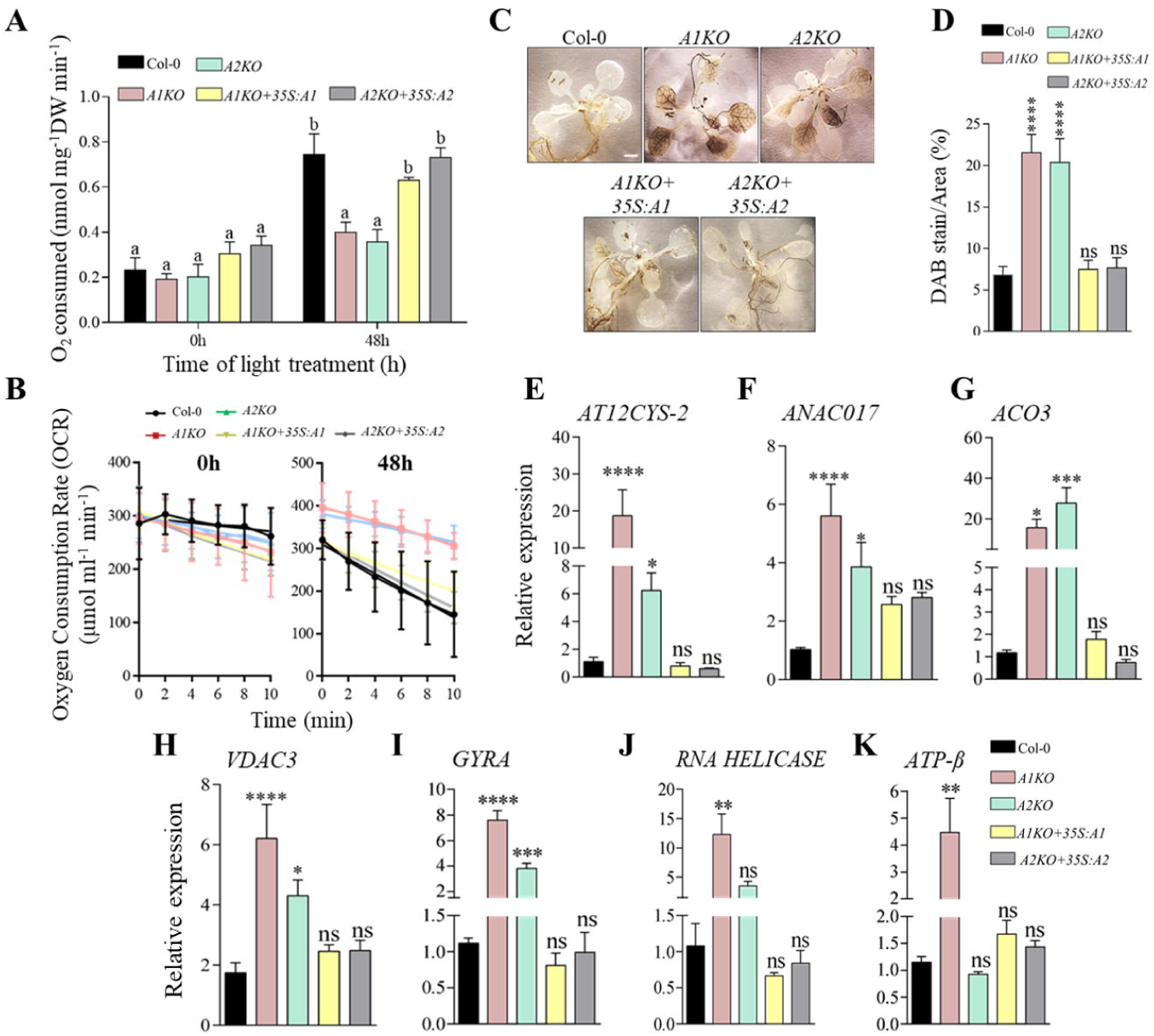
Loss of atDjA1 and atDjA2 impairs mitochondrial functions. **A)** Seeds of indicated genotypes were allowed to germinate for 0h and 48h before measuring the Oxygen consumption (nmol O_2_ mg^-1^ DW min^-1^) using a Clark-type O_2_ electrode at 25°C. Graphical depiction of the oxygen consumption per mg dry weight of seeds germinated in light for 0 and 48 hours. **B)** The rate graph depicts the rate of oxygen consumption by seeds of different genotypes over time after 0h and 48h of light treatment. The results reflect the mean ± SE of three independent studies. Columns with different letters indicate significant differences (Tukey’s test, Two-way ANOVA). **C)** Three-week-old seedlings of the indicated genotypes grown on ½ MS were stained with DAB to detect H_2_O_2_ observed as dark brown stain. **D)** Graphical representation of the DAB staining intensity of indicated genotypes measured using ImageJ. Error bars indicate SE of means of three independent studies. Asterisks denote statistically significant differences (****, *P<0.0001*, Dunnett’s test One-way ANOVA) ns – not statistically significant (P>0.05). **E-K)** Expression levels of the indicated genes, **E)** *AT12CYS-2* (regulator of MRS, AT5G09570), **F)** *ANAC017* (regulator of MRS, AT1G34190), **G)** *ACO3* (Mitochondrial dysfunction stimulon/MDS component, AT2G05710), **H)** *VDAC3* (MDS component, AT5G15090), **I)** *GYRA* and **J)** *RNA HELICASE* (mitochondrial biogenesis; AT3G10690 and AT3G26560, respectively) and **K)** *ATP-*β (respiratory complex component, AT5G08690) were analyzed by qRT-PCR in three-week-old seedlings of the indicated genotypes. The fold change was normalized against Arabidopsis *ACTIN* and *GAPDH or EL*α*1* genes. The results reflect the mean ± SEM of three independent experiments. Asterisks (****, *P<0.0001*; ***, *P<0.001*; **, P*<0.01*; and *, *P<0.05*, Dunnett’s test, One-way ANOVA) indicate significant differences. ns – not statistically significant (*P>0.05)*.

### Knockout of atDjA1 and atDjA2 results in increased ROS accumulation

An important feature of mitochondrial respiration is the production of ROS like superoxides (O_2_^-^) and subsequently hydrogen peroxide (H_2_O_2_). Defects in mitochondrial respiration frequently result in excessive ROS generation in complexes I and III of the ETC (Zorov, Juhaszova and Sollott, 2014). Controlled production of ROS acts as a signal in various pathways, however, uncontrolled production of ROS leads to oxidative damage to the cellular components. The equilibrium between reactive oxygen species (ROS) that are regularly produced in the process of respiration and scavenging of ROS ultimately determines the amount of hydrogen peroxide (H_2_O_2_) that is released from mitochondria, which act as an important signal molecule (Aon *et al*., 2012). To determine if anomalies in mitochondrial respiration in atDjA1 and atDjA2 mutants conduce to ROS accumulation, *A1KO* and *A2KO* seedlings were stained with DAB histochemical dye for the presence of H_2_O_2_, a major ROS marker **(Fig. 4C)**. A significant increase of about ten-fold in staining intensity was observed in both *A1KO* and *A2KO* leaves when compared to Col-0 (*P<0.0001*) under non-stressed control circumstances **(Fig. 4, C and D)**. As expected, ROS levels in the complemented lines *A1KO+35S:A1* and *A2KO+35S:A2*, was comparable to Col-0 **(Fig. 4, C and D)**, establishing a correlation between the elevated ROS accumulation and the respiratory defects in atDjA1 and atDjA2 knockout lines.

### Knockout of atDjA1 and atDjA2 activates Mitochondrial Retrograde Signaling (MRS)

Above, we show that downregulation of atDjA1 and atDjA2 impairs seed germination, most probably because of mitochondrial dysfunction. Several nuclear genes are known to be triggered in response to ROS buildup induced by inhibitors of mitochondrial respiration or constitutive mitochondrial malfunction caused by genetic abnormalities - a process known as Mitochondrial Retrograde Signaling (MRS) (Clifton *et al*., 2005; Schwarzländer *et al*., 2011; He *et al*., 2022). qRT-PCR was used to examine if reduced respiration combined with an increase in ROS accumulation had resulted in MRS activation in *A1KO* and *A2KO* plants. A few genes were chosen that have previously been shown to be upregulated in response to mitochondrial dysfunction: *AT12CYS-2* and *ANAC017 -* regulators of MRS (Van Aken and Whelan, 2012; Wang *et al*., 2020); *ACO3* and *VDAC3* - Mitochondrial Dysfunction Stimulon (MDS) components (Pascual *et al*., 2021); *ATP-*β - an ETC component (Giegé *et al*., 2005; Busi *et al*., 2011) and *GYRA* and *RNA HELICASE* - components that aid in mitochondrial biogenesis (Wang *et al*., 2014; Tamadaddi *et al*., 2021) **(Fig. 4, E to K)**. In both *A1KO* and *A2KO* lines, the expression of MRS regulators *AT12CYS-2* and *ANAC017* was substantially increased, demonstrating the activation of MRS in the mutants **(Fig. 4, E and F)**. While *A2KO* also demonstrated a significant increase (6-8 folds in *AT12CYS-2* and 4-5 folds in *ANAC017*), *A1KO* revealed a much higher expression of both MRS regulators (20-25 folds in *AT12CYS-2* and 6-7 folds in *ANAC017*). *ACO3* and *VDAC3*, two components of MDS, was also upregulated in both *A1KO* and *A2KO* suggestive of mitochondrial defects **(Fig. 4, G and H)**. *ACO3* expression was higher (30-40 folds) in *A2KO* **(Fig. 4G)**, while *A1KO* displayed higher expression of *VDAC3* (6-7 folds) **(Fig. 4H)**. In *A1KO*, the expression of *GYRA* was significantly upregulated by approximately sevenfolds and that of *RNA HELICASE* by approximately tenfold. Conversely, *GYRA* expression in *A2KO* was four to five times greater than that of Col-0, although RNA helicase expression remained comparable **(Fig. 4, I and J)**. *ATP-*β additionally displayed a fourfold increased expression in the absence of atDjA1, but was unaffected in *A2KO* seedlings **(Fig. 4K)**. The complemented lines *A1KO+35S:A1* and *A2KO+35S:A2* exhibited expressions similar to Col-0 for all the genes examined, thereby demonstrating that compromised mitochondrial homeostasis in atDjA1 or atDjA2 mutants activates MRS **(Fig. 4, E to K)**.

### Loss of atDjA1 reduces the number of active mitochondria per cell

The loss of mitochondrial import-associated proteins, as well as faulty import of various other nuclear-encoded mitochondrial proteins, have been shown to decrease the number of mitochondria per cell (Hye *et al*., 2004; Wall, Mitchenall and Maxwell, 2004; Tamadaddi *et al*., 2021). Because atDjA1 and atDjA2 are likely involved in the import of mitochondrial proteins from the cytosol, it was hypothesized that their decreased levels might impact the number of mitochondria per cell. To investigate this, protoplasts of *A1KO*, *A2KO* and Col-0 plants were stained with MitoTracker Red, which exclusively stains active mitochondria. The number of mitochondria visible as red puncta was then counted in these protoplasts. Compared to Col-0, *A1KO* protoplasts exhibited substantially fewer mitochondria (about 50% less, *P<0.0001*), whereas *A2KO* protoplasts showed no difference **(Fig. 5, A and B)**. Mitochondrial number was restored in *A1KO+35S:A1* complemented lines to levels comparable to Col-0 **(Fig. 5B)**, strongly indicating a substantial dependence of Arabidopsis on atDjA1 for either mitochondrial biosynthesis or maintenance.

**Figure 5.**
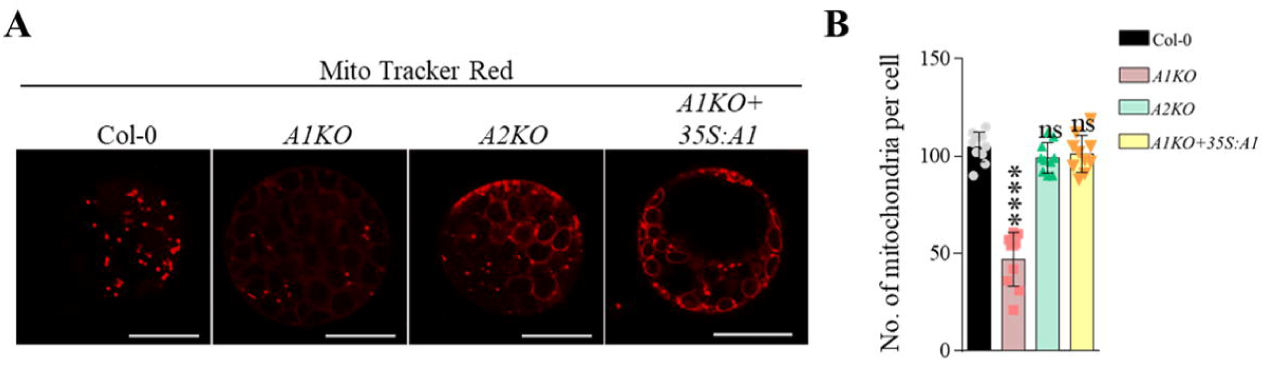
Loss of atDjA1 affects the number of functional mitochondria. **A)** Protoplasts obtained from three-week-old plants of indicated genotypes were stained with MitoTracker red (500 nM, 570-620 nm) and analyzed for the total number of active mitochondria using confocal microscope Mitochondria visible as red dots were counted using ImageJ’s “Analyze particle” tool. **B)** Graphical representation of the total number of active mitochondria in the protoplasts of the indicated genotypes. The results reflect the mean ± SE of a total of at least 15 distinct protoplasts of each genotype obtained from three independent preparations (from plants grown independently). Asterisks indicate significant differences (****, *P<0.0001*, Dunnett’s test, One-way ANOVA). ns – not statistically significant (*P>0.05*). Scale bar =10 µm.

### Knockout of atDjA1 affects mitochondrial protein import

There were fewer functioning mitochondria per cell in protoplasts isolated from atDjA1 knockout plants. So, we hypothesized that mitochondrial protein import would be impeded in *A1KO* lines. For this we employed a construct (mt gk/ *CD3 987*) that expresses GFP with an N-term mitochondrial targeting sequence (MTS). MTS-GFP is efficiently guided into the mitochondria where specific proteases cleave the MTS resulting in the expression of a processed GFP within the mitochondrial matrix (Köhler *et al*., 1997; Nelson, Cai and Nebenführ, 2007). To assess the import defect, the amount of GFP colocalizing with MitoTracker Red stained mitochondria was quantified in Col-0 and *A1KO* protoplasts. In the WT protoplasts, the majority of the GFP puncta colocalized with the mitochondria. However, as shown by the merged image, in the *A1KO* protoplasts there was a significant decrease in the colocalization of GFP signals with the mitochondria **(Fig. 6A)**. The Pearson coefficient of localization (r) was calculated from the microscope images using Image J software to quantify the same. The analysis showed a 50% reduction in the colocalization of GFP and MitoTracker Red signals in *A1KO* compared to Col-0 (*P=0.005*) **(Fig. 6B)**. Accumulation of a premature band of MTS-GFP (M) in *A1KO* samples as compared to Col-0 further validates our microscopy data **(Fig. 6C)**. These results thus point to a major involvement for the cytosolic Class I JDP atDjA1 in the mitochondrial import of cytosolic proteins.

**Figure 6.**
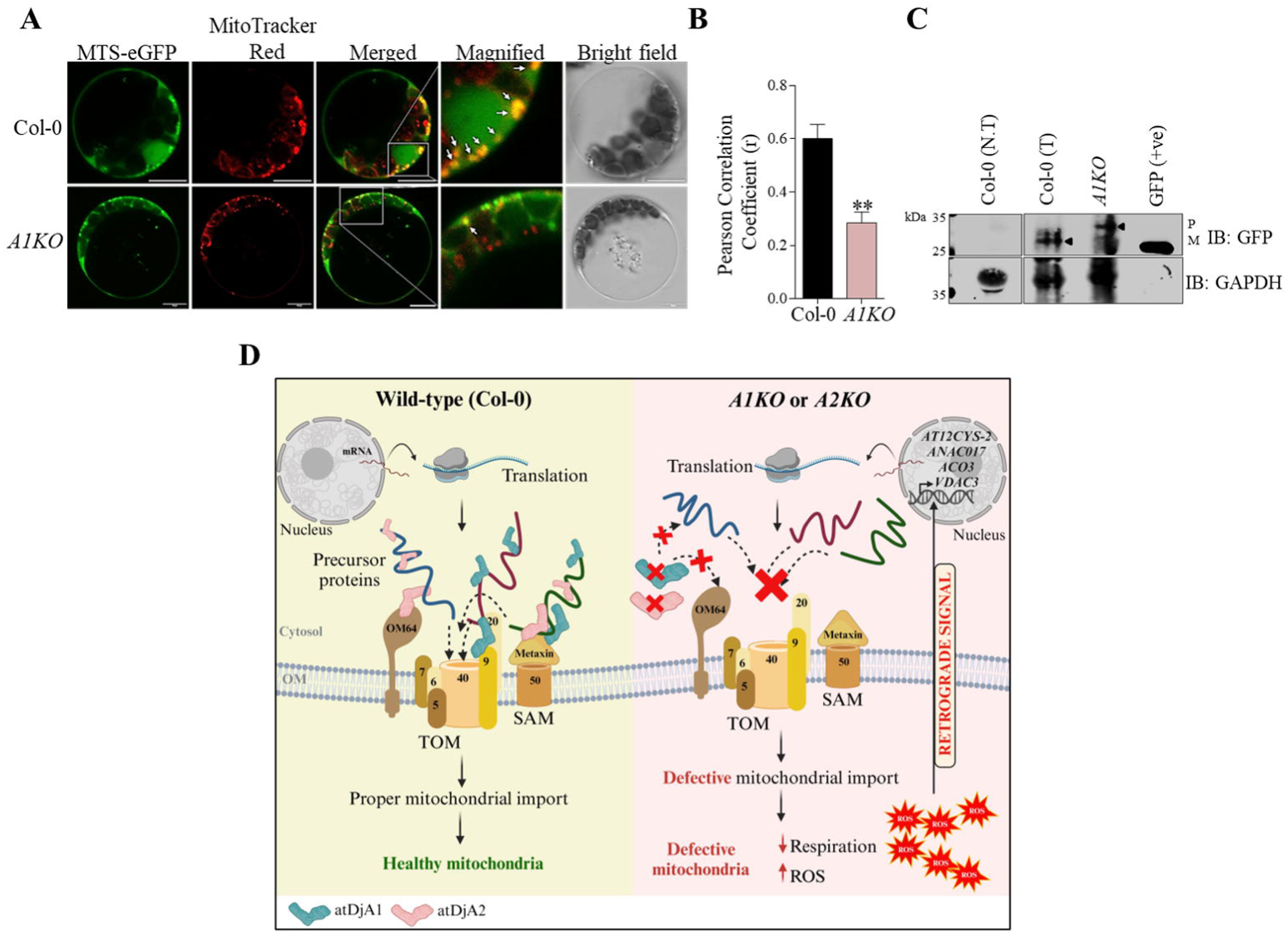
Loss of atDjA1 results in mitochondrial import defect in *A. thaliana.* **A)** Protoplasts isolated from indicated genotypes were transfected with the construct encoding fusion protein, MTS-GFP (mt-gk/*CD3 987*) and visualized using a confocal microscope after 20h of transfection. The colors green, red, and blue correspond to the representation of GFP (488-520 nm), mitochondria stained with MitoTracker red (570-620 nm) and chlorophyll autofluorescence (650-750 nm), respectively. Scale bar = 10 µm. **B)** Quantification of colocalization of MTS-eGFP with MitoTracker Red by Pearson correlation coefficient (r); each bar represents the average correlation coefficient (±SD) of at least nine different protoplasts from three biological replicates (**, *P=0.005,* Student t-test). **C)** Equal amounts of protein extracts isolated from protoplasts of specified genotypes transfected with MTS-GFP were resolved by SDS-PAGE, electroblotted onto nitrocellulose membrane and probed with anti-GFP antibody to identify the cytosolic precursor form (P) and mature mitochondrial form (M) of GFP protein. GAPDH was employed as the negative control. N.T-non-transfected, T-transfected sample **D)** Possible working model of how *A. thaliana* cytosolic Class I JDPs atDjA1 and atDjA2 help in preserving mitochondrial homeostasis. atDjA1 and atDjA2 are cytosolic JDPs, which bind to mitochondrial precursor proteins in the cytosol, thus keeping them in an unfolded, import-competent confirmation while bringing them to the mitochondria for import. Through their interactions with selected mitochondria outer membrane (OM) receptors, atDjA1 and atDjA2 help in proper import of pre-proteins and contribute to the healthy functioning of mitochondria. Absence of atDjA1 or atDjA2 causes defects in import of precursor proteins, resulting in decreased mitochondrial respiration and an increased production of mitochondrial ROS. The increased ROS act as signaling molecules for activating mitochondrial retrograde signaling, which modulates the expression of several nuclear genes.

## Discussion

Class I JDPs are inarguably the most conserved and ubiquitous of all JDPs. In addition to being important for the maintenance of cellular PQC, cytosolic Class I JDPs such as DnaJA1, DnaJA2 and DnaJA4 in *H. sapiens* and Ydj1 and Xdj1 in *S. cerevisiae* also play an integral part in the translocation of proteins from the cytosol to other cellular compartments such as the mitochondria and ER (Caplan, Cyr and Douglas, 1992; Kanazawa *et al*., 1997; Bhangoo *et al*., 2007; Sahi *et al*., 2013). Arabidopsis cytosol harbors two such proteins, atDjA1 and atDjA2 (Verma *et al*., 2017) and in this study we explored their role in mitochondrial protein import and homeostasis.

In budding yeast, cytosolic Class I JDP, Xdj1 associates with outer mitochondrial surface receptors to facilitate protein import (Opaliński *et al*., 2018). Although both atDjA1 and atDjA2 complement the slow growth phenotype of Xdj1 deletion, atDjA2 imports Mdj1 into the mitochondria more efficiently. This ability of atDjA2 to import Mdj1 into the yeast mitochondria is lost when specific residues in the CTD are substituted to the corresponding residues present in atDjA1, indicating that the differences in mitochondrial import function of these, almost identical JDPs stem from subtle variations in their client binding domains. This was further unveiled in our pulldown assays. atDjA1 interacted with Tom9.2, and atDjA2 interacted with OM64. Tom9.2 is involved in the import of Tom40 and the assembly of the TOM complex and OM64 is associated with the import of respiratory complex precursor proteins such as the F_A_D component of the ATP synthase (Lister *et al*., 2007; Parvin *et al*., 2017). Therefore, atDjA1 and atDjA2 could be conveying specific precursor proteins from the cytosol to Tom9.2 and OM64, respectively, to facilitate their import into the mitochondria **(Fig. 6D)**. Although speculative at the moment, this indicates the ability of atDjA1 and atDjA2 to carry out unique mitochondrial import functions in Arabidopsis. This is analogous to the role of cytosolic Class I JDPs in mitochondrial import in budding yeast. While both Xdj1 and Ydj1 are involved in the import of proteins into mitochondria in *S. cerevisiae*, Xdj1 has been explicitly assigned for this task (Sahi *et al*., 2013). Xdj1 and not Ydj1, interacts with the outer mitochondrial receptor protein Tom22 *via* its client binding domain (CTD) to promote the import of numerous precursor proteins into mitochondria (Opaliński *et al*., 2018).

The mitochondria in dry seeds are inactive due to the absence of cristae and respiratory complexes (Law *et al*., 2012). Proper germination requires the timely activation of these quiescent mitochondria, which in turn depends on proper import of a multitude of proteins which subsequently get sorted to different mitochondrial subcompartments (Farooq *et al*., 2021). Our findings reveal that atDjA1 and atDjA2 are crucial for seed germination, cotyledon opening and seedling establishment, especially under stress conditions. While our results are consistent with the involvement of atDjA1/A2 in mitochondrial homeostasis, we cannot completely rule out the possibility that other cytosolic PQC activities are also disrupted in these mutants. For example, Ydj1 of *S. cerevisiae* plays a crucial role in cellular stress response and PQC, including protein folding and refolding, inhibition of protein aggregation, dissolution of stress granules (Cry, 1995; Lu and Cyr, 1998; Walters *et al*., 2015). Additionally, Ydj1 safeguards newly formed protein kinases and regulates the rate of their maturation under stressful circumstances (Mandal *et al*., 2008). In a similar vein, atDjA1 and atDjA2 might be engaged in diverse PQC activities in the cytosol. Although not much is known about atDjA2, atDjA1 has been shown to regulate intracellular pH levels and affect root development under high salt concentrations (Yang *et al*., 2010). Further, loss of atDjA1 is also associated with abiotic stress hypersensitivity (Salas-muñoz *et al*., 2016). Nonetheless, since both atDjA1 and atDjA2 mutants showed poor respiration and increased ROS buildup, we propose that atDjA1/A2 are crucial for preserving mitochondrial homeostasis in Arabidopsis. Impaired mitochondrial protein import hampers the availability of essential components involved in respiration, ROS regulation, mtDNA maintenance, and numerous other vital biochemical activities (Millar *et al*., 2008; Kawamata and Manfredi, 2010; Ghifari and Murcha, 2022). This explains the delayed seed germination, respiratory defects and ROS accumulation in atDjA1 and atDjA2 mutants.

Reduced number of functional mitochondria in atDjA1 mutants may result from either defects in mitochondrial biogenesis or the presence of mitochondria with compromised membrane potential, both of which could result from deficient protein import from the cytosol (Franco-Iborra *et al*., 2018; Maruszczak, Ayyamperumal and Chacinska, 2023). While mitochondrial number was not compromised in atDjA2 mutant, activation of MRS in both atDjA1 and atDjA2 mutants, does underscore the fact that both these JDPs are required for maintaining optimum mitochondrial function in Arabidopsis (Schwarzländer *et al*., 2011; Wang *et al*., 2016; He *et al*., 2022). An upsurge in the expression of *VDAC3*, *ACO3*, *ATP-*β, *GYRA* and *RNA HELICASE* genes, which are important for metabolite transport, biochemical reactions, respiration and maintenance preservation of mitochondrial DNA, may serve as a compensatory mechanism to deal with mitochondrial stress or dysfunction in these mutants **(Fig. 6D)**.

In summary, our results enhance our understanding of cytosolic Class I JDPs in *A. thaliana*. Since atDjA1 and atDjA2 are very similar proteins, functional redundancy with respect to maintaining mitochondrial homeostasis is not surprising. However, atDjA1 mutant exhibited more severe phenotypes compared to that of atDjA2. While functional differences between atDjA1 and atDjA2 can be attributed to their unique clientele on the mitochondrial surface, differences in their expression profiles may also be contributing to their functional specificities in Arabidopsis. We show that atDjA1 is at least two-fold more abundant than atDjA2. The observed defects in atDjA2 mutants, despite the presence of a seemingly identical, highly abundant atDjA1 paralog, strongly correspond to their distinct, yet unknown functionalities in mitochondrial protein import in Arabidopsis.

## Materials and methods

### In-silico analysis

Multiple alignments of protein sequences were done using MAFFT software (https://www.ebi.ac.uk/Tools/msa/mafft/) (Katoh and Standley, 2013) and ESPript was employed to generate the alignment figure (https://espript.ibcp.fr/ESPript/cgi-bin/ESPript.cgi) (Gouet, Robert and Courcelle, 2003). The tertiary structures were predicted using the SWISS-MODEL web server (https://swissmodel.expasy.org/) (Waterhouse *et al*., 2018), after which they were processed using PyMol (DeLano, 2002). The graphs from various studies were plotted and statistically analyzed using the GraphPad Prism 6.0 software(https://www.graphpad.com/scientificsoftware/prism/www.graphpad.com/scientific-software/prism/) (Swift, 1997). Illustrations were created using the Bio Render software (https://www.biorender.com/) (Perkel, 2020).

### Plant methods and growth conditions

This study employed *Arabidopsis thaliana* plants, ecotype Columbia (Col-0). SALK T-DNA insertion mutant lines for atDjA1 (SALK_132923) and atDjA2 (SALK_071563), were acquired from the Arabidopsis Biological Resource Center (ABRC, Ohio State University, USA). After surface sterilization, seeds were placed on ½ MS plates (0.6 percent (w/v) agar), with or without sucrose. Following stratification in the dark at 4°C for 48h, the plates were moved to a Percival LED22C8 growth chamber at 22°C Day/18°C Night and 70% humidity [light intensity 100±20 μmol m^-2^ s^-1^, 16h light:8h dark cycle]. Agrobacterium (GV3101)-mediated floral dip transformation, as reported in Zhang *et al*., 2006, was used to create atDjA1/A2 complementation lines (referred to as *A1KO+35S:A1* and *A2KO+35S:A2* hereafter). Equally aged seeds from wild type, mutant and complementation lines were used for phenotypic assays. Seeds were plated on ½ MS plates with varying concentrations of osmotic and ionic stressors such as mannitol, glucose and NaCl for phenotypic studies. The experimental plates for glucose-based assays had different concentrations of glucose and no sucrose. The phenotypic experiments in the presence of abiotic stressors were conducted for seven days (168h) to examine radicle emergence and open cotyledons.

### Plasmid construction for yeast experiments, complementation and sub-cellular localization

Full-length ORFs of atDjA1 and atDjA2 were PCR amplified using primers with suitable restriction sites from cDNA prepared from wild type Arabidopsis seedlings. Site-directed mutagenesis was done by Quick change mutagenesis method as per manufacturer’s protocol (QuikChange ® Site-Directed Mutagenesis Kit). For protein expression, full-length ORFs of atDjA1 and atDjA2 were cloned in BamH1/Xho1 site in pSMT3 vector (kind gift from Christopher D. Lima, Sloan Kettering Institute, USA) with an N-terminal His+SUMO tag. For affinity pull-down experiments, coding sequences corresponding to the cytosolic domains of *A. thaliana* outer mitochondrial membrane proteins - Tom9.2 (aa 1-53, BamH1/Sal1), Metaxin (aa 1-191, BamH1/EcoR1) and OM64 (aa 488-586, BamH1/EcoR1) were PCR amplified from total Arabidopsis cDNA and cloned in pGEX-6p1 vector with an N-terminal GST tag. Clones for sub-cellular localization and mutant complementation were generated using Gateway® Technology as per the manufacturer’s instructions (Invitrogen). The sequences of interest were amplified from cDNA employing primers with attB1 and attB2 sites that flank gene-specific nucleotides, then cloned in pDONR207 (Invitrogen) using the BP reaction to create entry clones, and subsequently shuttled to the appropriate destination vectors via the LR reaction. For subcellular localization, full-length cDNA sequences of atDjA1 and atDjA2 with stop codons were shuttled to the pGWB606 destination vector with a 35S promoter for N-terminal GFP tagging (Nakagawa *et al*., 2007). For making complementation lines, the full-length cDNA sequences of atDjA1 and atDjA2 with stop codons were shuttled to the HBP047 destination vector (modified pCAMBIA1300 vector containing the gateway cassette) to drive the expression under the CaMV-35S promoter. All the clones generated were confirmed by sequencing. All primers used for cloning are listed in **Supplemental Table S1.**

### RNA isolation, cDNA synthesis and qRT-PCR

Total RNA was extracted from different plant tissues using TRIzol™ Reagent (Invitrogen) as described previously with minor modifications (Chomczynski and Mackey, 1995). Briefly, the homogenate obtained from 100-150 mg of tissue was vigorously shaken with chloroform and centrifuged for 5 min at 10,000 xg at 4°C. The upper aqueous-phase was mixed with 1 volume of isopropyl alcohol and incubated at −80°C for 30 min, followed by centrifugation. The resulting RNA pellet was washed with 75% ethanol, air-dried, dissolved in DEPC-treated water, and subsequently treated with DNaseI (Invitrogen). The quality of RNA was assessed by gel electrophoresis and quantified using an Eon Microplate Spectrophotometer (BioTek). iScript cDNA synthesis kit (Bio-Rad) was employed to reverse-transcribe 2 µg of the DNase I-treated RNA, after which the cDNA was diluted to a final concentration of 40 ng/µl for further experiments. qPCR Master Mix (KAPA SYBR® FAST) was used for performing real-time PCR on the CFX CONNECT TM Real-Time PCR Detection System (Bio-Rad). *ACTIN2* and *ELF*α or *GAPDH* were used as internal controls. MIQE guidelines were followed for all data processing (Bustin *et al*., 2009).

### Protoplast isolation and transfection

Protoplasts were extracted from three-week-old seedlings, as previously reported (Yoo, Cho and Sheen, 2007). Briefly, the leaves were sliced into thin strips using a sharp razor and immediately placed into the enzyme solution (20 mM MES (pH 5.7), 1.5% (w/v) cellulase R10, 0.4% (w/v) macerozyme R10, 0.4 M mannitol, and 20 mM KCl) in a six-well plate, coated with BSA (5% w/v). Following vacuum treatment, the plate was kept undisturbed in dark for 5–6h to allow for the release of protoplasts. After neutralizing with W5 solution (2 mM MES (pH 5.7), 154 mM NaCl, 125 mM CaCl_2_, and 5 mM KCl), the tissue debris from the solution was removed by filtration. The protoplasts were spun down at 100 xg and resuspended in W5 solution and allowed to settle down on ice. Settled protoplasts were resuspended in MMG solution (4 mM MES (pH 5.7), 0.4 M mannitol and 15 mM MgCl_2_) and again allowed to settle at RT. The resulting protoplasts were mixed with plasmid constructs (10–15 µg in 10 µl) in an MCT, followed by PEG-CaCl_2_ solution (40% PEG 4000, 0.2 M mannitol and 100 mM CaCl_2_) for transfection. Following transfection, protoplasts were resuspended in W1 solution (4 mM MES (pH 5.7), 0.5 M mannitol and 20 mM KCl) and incubated for 12-16h in dark at RT before visualization. For visualising mitochondria, the protoplasts were incubated with MitoTracker Red CMXRos (500 mM, Invitrogen) at RT. The confocal microscope (LSM 780; Carl Zeiss) was employed to visualize the GFP (488/520 nm), mitochondria (570/620 nm) and chlorophyll autofluorescence (650/750 nm).

### Protein extraction from *A. thaliana* protoplasts

Protein extraction from protoplasts was carried out as described previously (Lee *et al*., 2019). Briefly, the transfected protoplasts, after incubation for the required time, were spun down and resuspended in 100 µl protoplast lysis buffer (50 mM Tris-HCl (pH 7.5), 150 mM NaCl, 1 mM EDTA, 1% Triton X-100, 1X Protease Inhibitor Cocktail (PIC), 10% glycerol), followed by brief sonication (Sonics USA). The solution was then clarified by centrifugation at 3000 xg for 10 min, and the supernatant was transferred to a fresh tube. 40 µg of total protein was mixed with SDS sample buffer (62.5 mM Tris–HCl (pH 6.8), 5% glycerol, 2% SDS, 2% β-mercaptoethanol, and 0.01% bromophenol blue) and resolved by SDS-PAGE. Western analysis was performed with anti-GFP (ABCAM) primary antibody, followed by Invitrogen’s Goat anti-rabbit secondary antibody.

### DAB staining for detection of H_2_O_2_

H_2_O_2_ was detected by 3,3’diaminobenzidine (DAB) staining as described previously with minor modifications (Thordal-Christensen *et al*., 1997). Three-week-old seedlings were immersed in freshly made DAB solution (1 mg/ml in H_2_O), vacuum infiltrated for 10 min, and incubated overnight at RT in the dark. The stained seedlings were bleached in acetic acid/glycerol/ethanol (1/1/3, v/v/v) solution at 95°C and stored in glycerol/ethanol (1/4, v/v) solution until the photographs were taken. Images were obtained using a Leica M205FA microscope. Quantification of staining intensity was done from the microscopy images using the ImageJ digital image analysis package.

### Seed respiration assay

Oxygen consumption by Arabidopsis seeds was evaluated using a Clark-type O_2_ electrode (Oxytherm, Hansatech Ltd., Norfolk, UK), as reported by Heidorn-Czarna *et al*., 2018. The experiment was conducted in 50 mM HEPES (pH 7.2). Specifically, 20 mg of dry seeds were surface sterilized and stratified for 48h. Following this, the seeds were either utilized immediately (0h) or kept under light for 48h for germination, before being moved to the chamber for measuring oxygen absorption. The oxygen measurement was conducted for a duration of 10 min for each sample.

### Yeast methods

Yeast transformations were done using the LiAc/SS carrier DNA/PEG method (Gietz and Schiestl, 2007). Overnight grown yeast culture was centrifuged and resuspended in LiAc-TE, and kept at 30°C to make competent yeast cells. 20 μl of competent cells were mixed with ∼500 ng of plasmid DNA, denatured ssDNA (single strand DNA, 2 mg/ml) and PEG-LiAc-TE solution (40% PEG, 0.1 M LiAc, 10 mM Tris-HCl (pH 8.0), 1 mM EDTA). This transformation reaction was kept at 30°C followed by heat shock at 42°C for 20 min. The cells were centrifuged, plated on appropriate dropout selection media and incubated at appropriate temperature.

Total protein was extracted from yeast cells following the protocol described earlier (Tak *et al*., 2023). Briefly, 0.7-1.0 OD_600_ equivalent cells were spun down, treated with 0.1 N NaOH and resuspended in SDS sample buffer. The proteins were resolved by SDS-PAGE, electroblotted onto nitrocellulose membrane (Bio-Rad, USA) and probed with appropriate antibodies (Mdj1 - a generous gift from Dr. Elizabeth Craig, UW-Madison); atDjA1 - produced in-house; GAPDH or GAPC1/2 (Agrisera). Invitrogen’s Goat anti-rabbit or anti-mouse (IgG (H+L) DyLight 680 or 800) secondary antibodies were used at a 1:20000 dilution. Blots were imaged by the Odyssey Classic Infrared Imaging System (LI-COR Biosciences).

### Protein purification, quantification, antibody production and affinity pull-down experiments

A single colony of *E. coli* Rosetta cells transformed with the desired protein expression clone was grown at 37°C overnight with continuous shaking until its OD_600_ reached 0.6-0.8. The culture was induced by 0.5 mM IPTG at 16°C overnight. The culture was centrifuged and resuspended in lysis buffer (His-tagged proteins: 200 mM Tris-HCl (pH 7.5), 500 mM NaCl, 20 mM Imidazole, 1 mM DTT, 1 mM PMSF; GST-tagged proteins: 50 mM Tris (pH 7.5), 150 mM NaCl, 10% Glycerol, 1X PIC and 1 mM DTT) with lysozyme (1 mg/ml) and incubated on ice for 30 min. Following sonication, the cell suspension was centrifuged. Bead binding was performed by adding washed beads (Ni-NTA agarose beads (Qiagen) for His-tagged proteins and GSH agarose beads (ABT) for GST-tagged proteins) into the supernatant at 4°C. The protein-bound beads were washed thoroughly with appropriate wash buffers (His-tagged proteins: thrice with the lysis buffer; GST-tagged proteins: lysis buffer with 300 mM, 500 mM and 700 mM NaCl). Proteins were eluted from the beads by adding appropriate elution buffer (His-tagged proteins: 20 mM Tris-HCl (pH 7.5), 50 mM NaCl, 300 mM Imidazole and 10% glycerol; GST-tagged proteins: 50 mM Tris-HCl (pH 7.5), 150 mM NaCl, 0.1 mM EDTA and 10 mM reduced glutathione) at 4°C. For generating antibodies, purified atDjA1 protein was dialyzed against 50 mM Tris-HCl (pH 8.0), 1 mM EDTA, 150 mM NaCl, 10% glycerol, and 3 mM β-ME, at 4°C. sENP protease was used in the dialysis membrane to remove His+SUMO tag. Cleaved atDjA1 protein was utilized for antibody generation at a local commercial facility (DPL, Bhopal, India). For GST pull-down assay, 5 µg of His+SUMO-atDjA1 or atDjA2 was incubated with GST-tagged Tom9.2, Metaxin or OM64 on GSH beads at 4°C for 1h as described earlier (Hosoda *et al*., 2003). Bead-bound GST tag or empty GSH beads were used as controls. Beads were thoroughly washed, and the proteins associated with the beads were subsequently resolved on SDS-PAGE before being detected by Western blot analysis with anti-GST (Invitrogen) and anti-atDjA1 antibodies.

### Accession numbers

The sequence information of the genes examined in this study was sourced from the Saccharomyces Genome Database (SGD) and The Arabidopsis Information Resource (TAIR) databases. The accession numbers for these genes are as follows: Xdj1 (YLR090W), atDjA1 (AT3G44110), atDjA2 (AT5G22060), Tom9.2 (AT5G43970), OM64 (AT5G09420), Metaxin (AT2G19080), *ANAC017* (AT1G34190), *ACO3* (AT2G05710), *VDAC3* (AT5G15090), *ATP-*β (AT5G08670), *GYRA* (AT3G10690), *RNA HELICASE* (AT3G26560), *AT12CYS-2* (AT5g09570), *ACTIN* (AT3G18780), *EL*α*1* (AT1G07940) and *GAPDH* (AT3G26650).

## Acknowledgements

The authors thank the Fund for Improvement of Science & Technology Infrastructure (FIST) for the live-cell imaging system. S.S.L., N. and Y.Y.R.P. thank IISER Bhopal, Council of Scientific and Industrial Research (CSIR) and Department of Biotechnology (DBT) respectively for the fellowship. The authors thank BioRender.com (https://biorender.com) for creating the illustrations.

## Author contributions

C.S. and A.K.V. conceived and initiated the project; C.S. supervised the research and furnished laboratory facilities and funding; S.S.L. and C.S. designed the experiments and wrote the manuscript; S.S.L. performed most of the experiments and analyzed the data; N. contributed to microscopic imaging and seed respiration assays and provided critical feedback on the manuscript; Y.Y.R.P. and S.P. generated resources and assisted with the experiments.

## Data availability

The authors confirm that all experimental data are available and accessible via the main text and/or the supplemental data.

