## Supplemental Figures for "Cytosolic Class I J-domain proteins aid mitochondrial protein import and influence homeostasis in *Arabidopsis thaliana*"

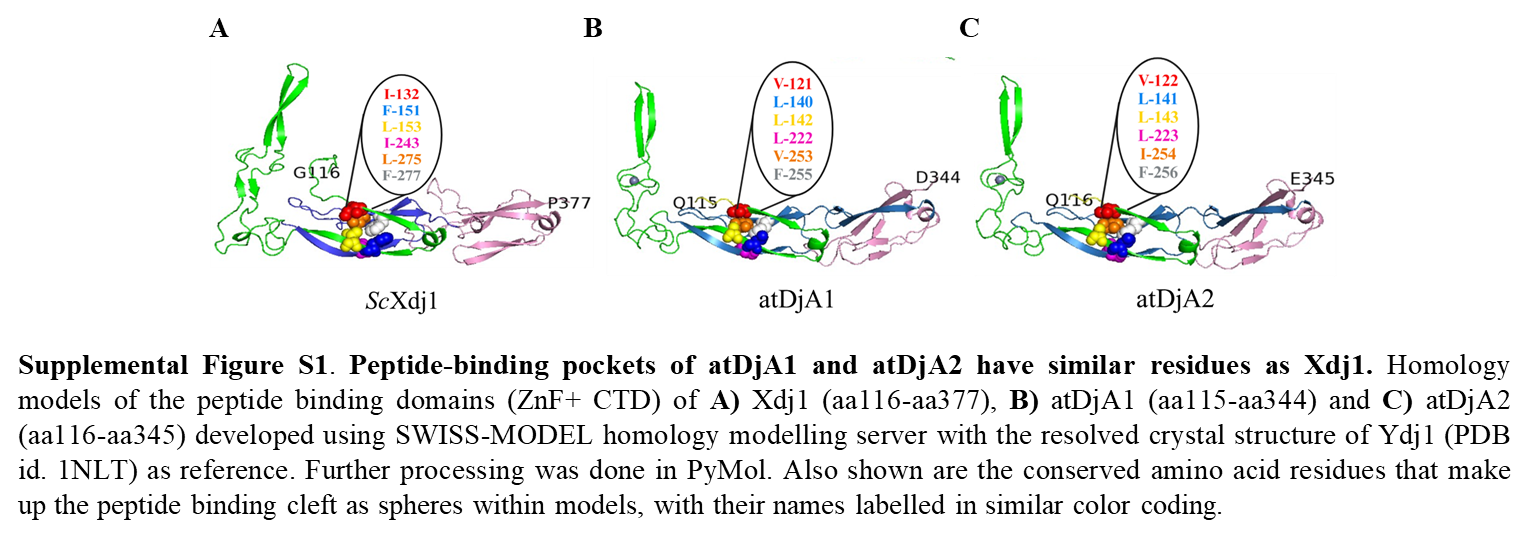

**Supplemental Figure S1**. **Peptide-binding pockets of atDjA1 and atDjA2 have similar residues as Xdj1.** Homology models of the peptide binding domains (ZnF+ CTD) of **A)** Xdj1 (aa116-aa377), **B)** atDjA1 (aa115-aa344) and **C)** atDjA2 (aa116-aa345) developed using SWISS-MODEL homology modelling server with the resolved crystal structure of Ydj1 (PDB id. 1NLT) as reference. Further processing was done in PyMol. Also shown are the conserved amino acid residues that make up the peptide binding cleft as spheres within models, with their names labelled in similar color coding.

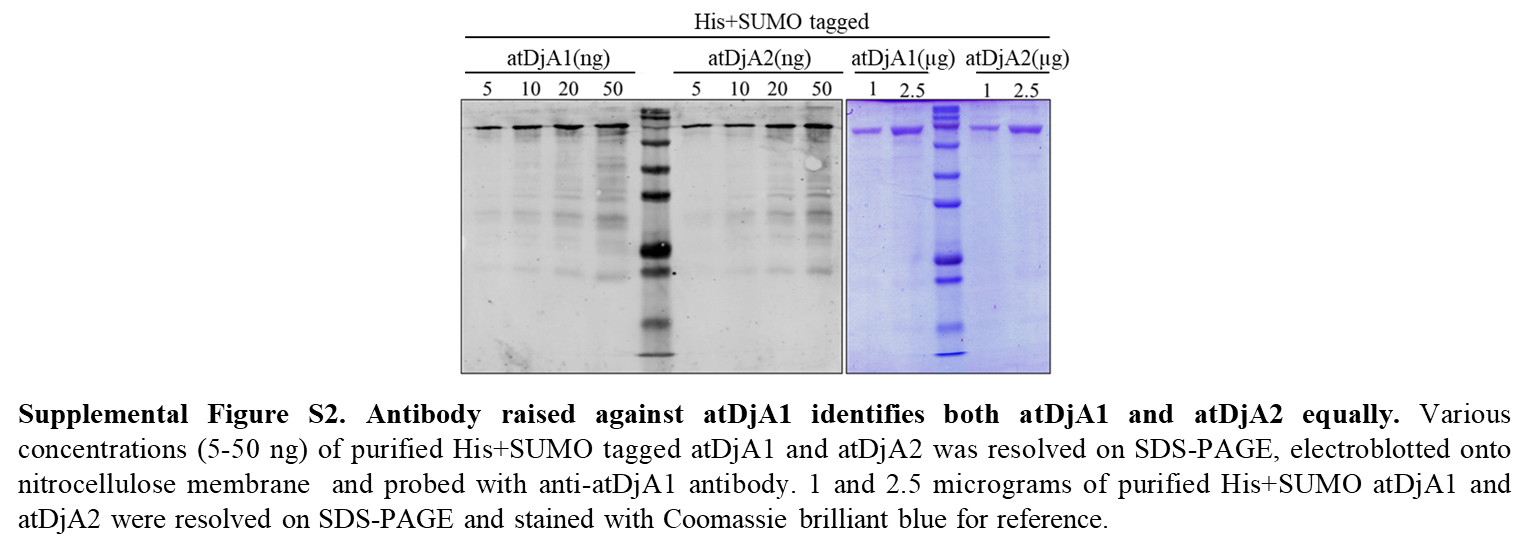

**Supplemental Figure S2. Antibody raised against atDjA1 identifies both atDjA1 and atDjA2 equally.** Various concentrations (5-50 ng) of purified His+SUMO tagged atDjA1 and atDjA2 was resolved on SDS-PAGE, electroblotted onto nitrocellulose membrane and probed with anti-atDjA1 antibody. 1 and 2.5 micrograms of purified His+SUMO atDjA1 and atDjA2 were resolved on SDS-PAGE and stained with Coomassie brilliant blue for reference.

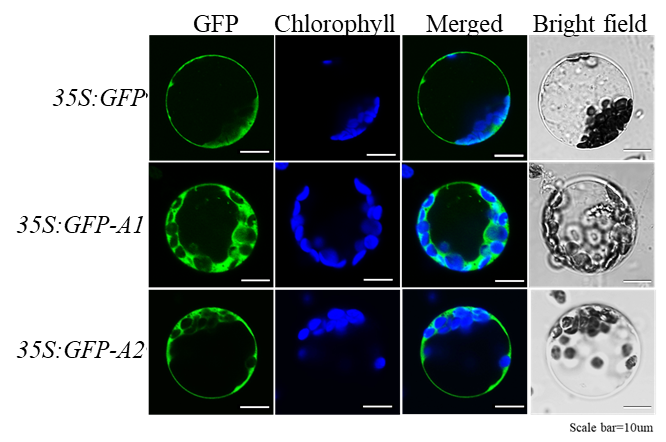

**Supplemental Figure S3. atDjA1 and atDjA2 are localized in the cytosol in *A. thaliana.*** Mesophyll cell protoplasts obtained from 3-weekold wild-type Arabidopsis plants were transfected with 35S:GFP-EV, 35S:GFP-atDjA1 and 35S:GFP-atDjA2 constructs. Following transfection, the cells were incubated for 12 hours before being examined using a live cell imaging microscope. Green fluorescent signals (488-520 nm), chlorophyll autofluorescence (blue, 650-750 nm), an overlay of GFP and chlorophyll signals and bright-field images are displayed. The scale bar is 10µm.

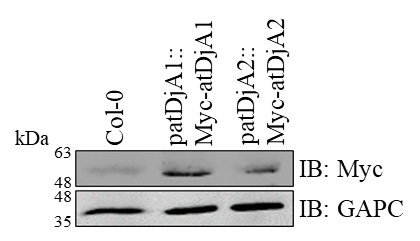

**Supplemental Figure S4. atDjA1 protein is abundant.** 0.5-0.7 g of fresh tissue from 15-day old seedlings of Col-0 and stable lines expressing N-term Myc-tagged atDjA1 and atDjA2 (under endogeneous promoter) were homogenized in 1 ml of Protein Isolation Buffer, PIB (25 mM HEPES, 200 mM NaCl, 0.5 mM EDTA, 0.1% v/v Triton X- 100, 5 mM Є-amino-n-Caproic acid, 1 mM Benzamidine) and were spun down for clarifying the lysate. After quantifying protein concentration of the lysate by Bradford assay, 5 µg of each sample was mixed with SDS sample buffer, resolved by SDS-PAGE, electro-blotted onto nitrocellulose membrane and probed with anti-Myc (SIGMA) antibody. GAPC was employed as the loading control.

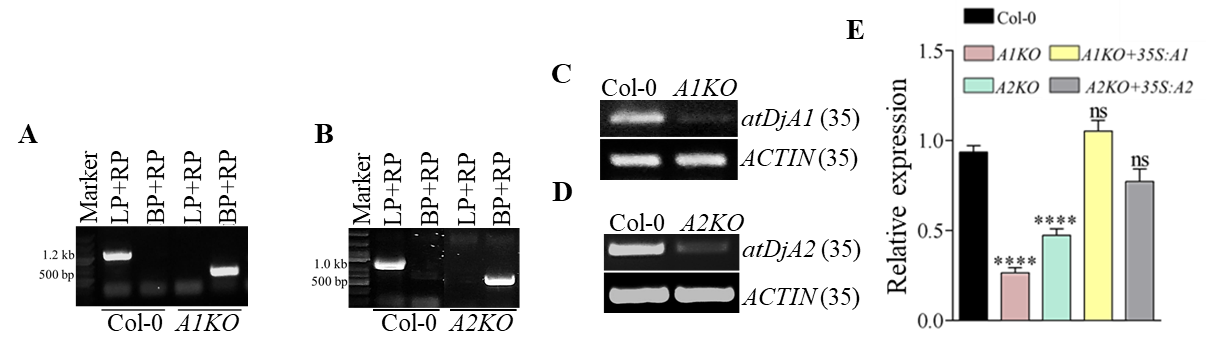

**Supplemental Figure S5. The loss of function mutants of atDjA1 and atDjA2 employed in phenotypic analysis.** Verification of the T-DNA insertion in **A)** atDjA1 and **B)** atDjA2 via genotyping of the SALK_132923 and SALK_071563 lines, respectively. SALK_132923 is denoted as *A1KO* and SALK_071563 is denoted as *A2KO*. Expression levels of **C)** atDjA1 and **D)** atDjA2 C-terminal fragments in T-DNA insertion lines by semi-quantitative RT-PCR. *ACTIN* gene was employed as internal control. PCR cycle number is indicated in brackets. **E)** Expression levels of atDjA1 and atDjA2 C-terminal fragments in T-DNA insertion lines and complemented lines by qRT-PCR analysis. The fold change was normalized against Arabidopsis *ACTIN2* and *ELα1* genes. Significant differences are indicated by asterisks (****, *P < 0.0001*, Dunnett's test, One-way ANOVA). The bars show the mean ± SE of three biological replicates. ns – not statistically significant (*P>0.05*).

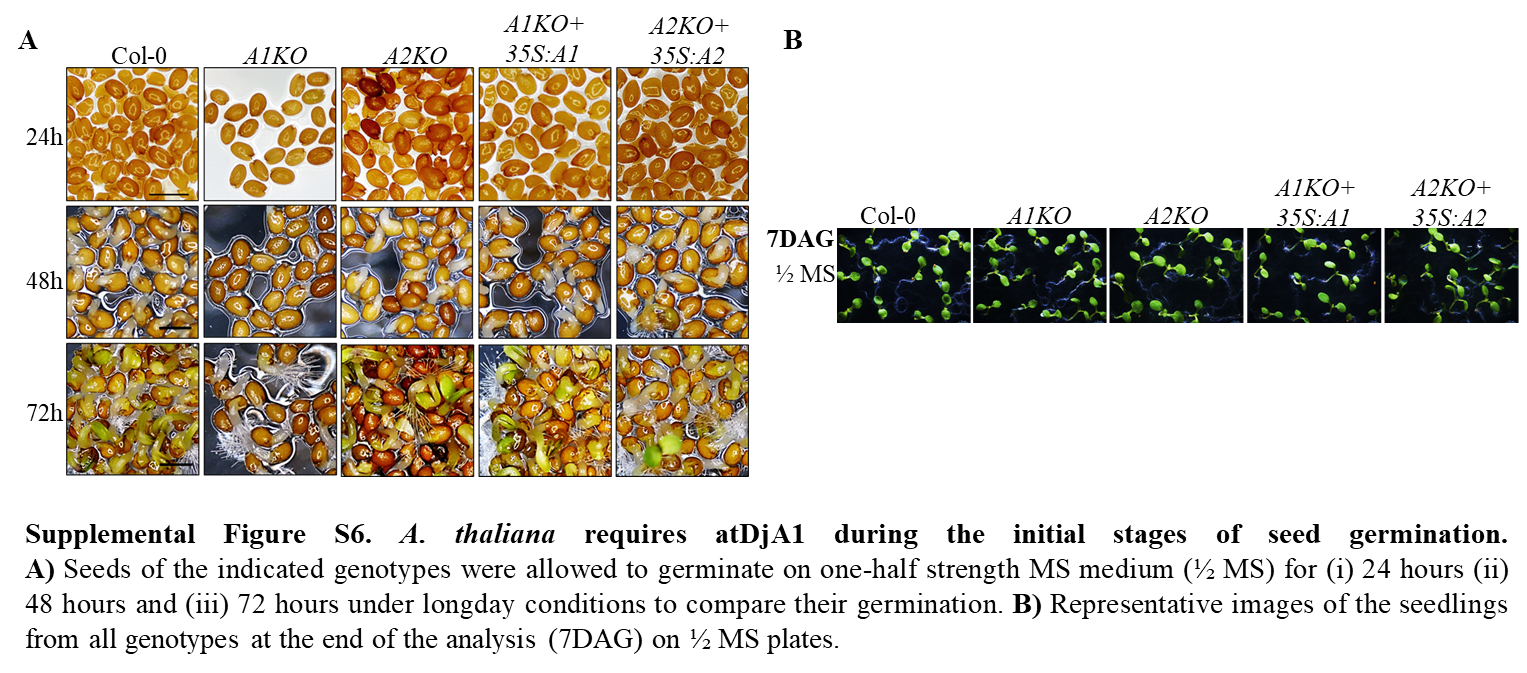

**Supplemental Figure S6.** ***A. thaliana* requires atDjA1 during the initial stages of seed germination.** **A)** Seeds of the indicated genotypes were allowed to germinate on one-half strength MS medium (½ MS) for (i) 24 hours (ii) 48 hours and (iii) 72 hours under longday conditions to compare their germination. **B)** Representative images of the seedlings from all genotypes at the end of the analysis (7DAG) on ½ MS plates.

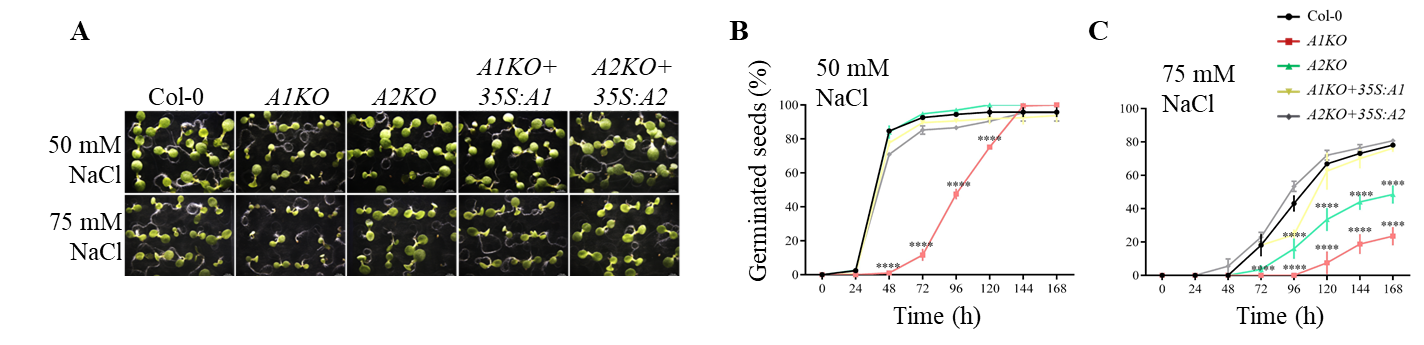

**Supplemental Figure S7. Germination of Arabidopsis under salt stress requires atDjA1 and atDjA2. A)** Seedlings of the indicated genotypes germinated for 7 days on 1/2 MS medium supplemented with 50 mM NaCl and 75 mM NaCl. **B-C)** Graphs displaying the germination rate of the specified genotypes during a 24-168-hour period on 50 mM NaCl (**B**) and 75 mM NaCl (**C**). The results are based on the mean ± SE of three biological replicates. An asterisk (****, *P<0.0001*, Tukey's test, One-way ANOVA) indicates a significant difference.

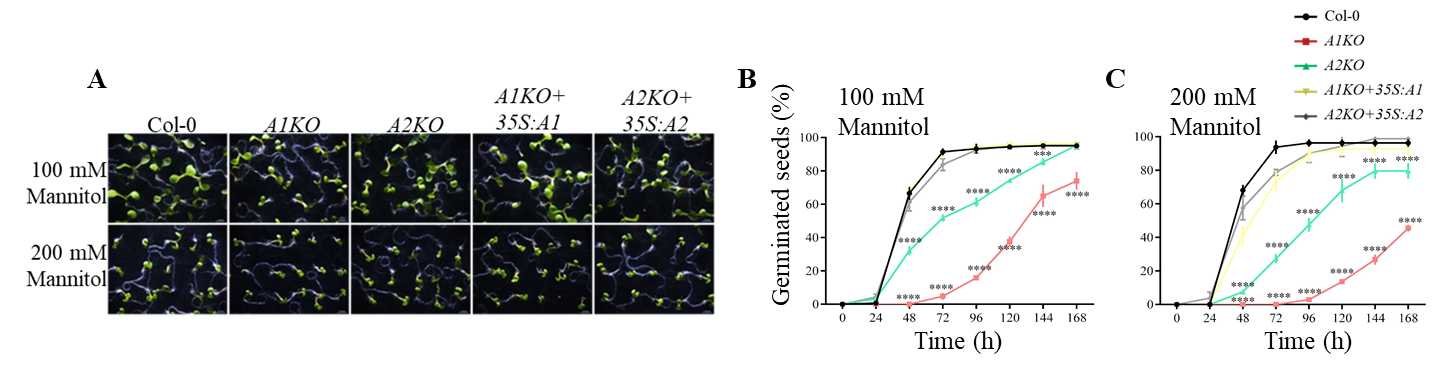

**Supplemental Figure S8. Germination of Arabidopsis under mannitol induced osmotic stress requires atDjA1 and atDjA2. A)** Seedlings of the indicated genotypes germinated for 7 days on 1/2 MS medium supplemented with 100 mM mannitol and 200 mM mannitol. **B-C)** Graphs displaying the germination rate of the specified genotypes during a 24-168-hour period on 100 mM mannitol (**B**) and 200 mM mannitol (**C**). The results are based on the mean ± SE of three biological replicates. An asterisk (****, *P<0.0001; ***, P<0.001*, Tukey's test, One-way ANOVA) indicates a significant difference.

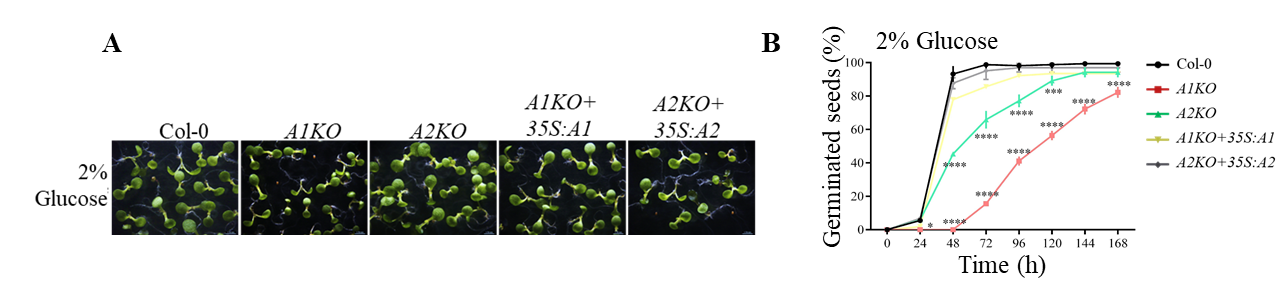

**Supplemental Figure S9. Germination of Arabidopsis under glucose induced osmotic stress requires atDjA1 and atDjA2. A)** Seedlings of the indicated genotypes germinated for 7 days on 1/2 MS medium supplemented with 2% glucose. **B)** Graph displaying the germination rate of the specified genotypes during a 24-168-hour period on 2% glucose. The results are based on the mean ± SE of three biological replicates. An asterisk (****, *P<0.0001; ***, P<0.001*; **, P<0.05*, Tukey's test, One-way ANOVA) indicates a significant difference.
