## Supplemental Table for "Cytosolic Class I J-domain proteins aid mitochondrial protein import and influence homeostasis in *Arabidopsis thaliana*"

**Supplemental material**

**Supplemental Table S1.** List of primers used in this study.

| **Yeast work-related** |
| --- |
| atDjA2_S218P_F      GAAGGTTGTCCCTGAGAAGAAGG  atDjA2_S218P_R      TCTCCTTTGCATTGTGGAC |
| atDjA2_K305N_F     CATCAAATCCAATCCCGGAGAGG  atDjA2_K305N_R     AGAAGCTGTCTTTTGTCC |
| **Protein work-related** |
| *At*Tom9.2_CD_F       CGCGGATCCATGGCGGCGAAGAGAATCGGAG      CCGGTAAATCTGGC  *At*Tom9.2_CD_R      ACGCGTCGACTTACGCTGCTTTTCCG |
| *At*Metaxin_CD_F     CGCGGATCCATGGAAGGCGATCAAGAGACG  *At*Metaxin_CD_R     CCGGAATTCTTACCTGTCTTCAAAGAGAAAC |
| *At*OM64_TPR_F      CGCGGATCCATGGCTTCTGAAGTTATG  *At*OM64_TPR_R      CCGGAATTCTTATTCAAGGACCAATGCG |
| atDjA1_F                  CGCGGATCCATGTTCGGTAGAGGACCCTC  atDjA1_R                  CCGCTCGAGTTACTGCTGGGCACATTG |
| atDjA2_F                  CGCGGATCCATGTTTGGAAGAGGACCTTC  atDjA2_R                  CCGCTCGAGTCACTGCTGGGCACATTG |
| **Genotyping and qRT-PCR-related** |
| atDjA1_LP_genotyping     TAGCAGCATTTGTTCTGCATG  atDjA1_RP_genotyping     TAATGAATGCAAAGGGACAGG |
| atDjA2_LP_genotyping     GGATTCTGATTGATAAAAGAAAAACC  atDjA2_RP_genotyping     TATGACTTCCGAATGGGTGTC |
| LB_6313R_SALK             TCAAACAGGATTTTCGCCTGCT |
| atDjA1_qRT_F_C-term     AGCTTTGTGTGGCTTCCAAT  atDjA1_qRT_R_C-term     GAAGTGGATGTAGAGCTTTACCC |
| atDjA2_qRT_F_C-term     CCTTGTGTGGCTTCCAGTTT  atDjA2_qRT_R_C-term     CGTGAAGTGAATGTATAGCTTACCC |
| atDjA1_F_FL             ATCCACTAGTATGTTCGGTAGAGGAC  atDjA1_R_FL            GCTTATCGATTTACTGCTGGGCACATTG |
| atDjA2_F_FL             ATCCACTAGTATGTTTGGAAGAGGACCTTC  atDjA2_R_FL             GCTTATCGATTCACTGCTGGGCACATTG |
| *ACTIN2.2*_qRT_F      GCCATCCAAGCTGTTCTCTC  *ACTIN2.2*_qRT_R      GCATGAGGAAGAGAGAAACCC |
| *ELα1*_qRT_F              AGGTTTTGAGGCTGGTATCTCT  *ELα1*_qRT_R              TGAAGACACCTCCTTGATGATTT |
| *GAPDH*_qRT_F         TTGGTGACAACAGGTCAAGCA  *GAPDH*_qRT_R        AAACTTGTCGCTCAATGCAATC |
| *AT12CYS-2*_qRT_F   TCCACGAACCTTTAAGCACG  *AT12CYS-2*_qRT_R   GCAGCCACCACGGAACC |
| *ANAC017_*qRT_F     CATCGGCTTTGTGGGCATTA  *ANAC017*_qRT_R     CACAAAGCACCCACGATTGA |
| *ACO3*_qRT_F           AAAATGACATTGTGGCGGCT  *ACO3*_qRT_R           AATTTCTTCCGTGGTTGGCC |
| *VDAC3*_qRT_F         ACAGCTACCCCGATTGTCAA  *VDAC3*_qRT_R        TGGTCGTGAAGTTGTGGCTA |
| *RNA HELICASE*_qRT_F   GAAGGCTAAAAACTTCTCAGGTCCATG  *RNA HELICASE*_qRT_R   CCACCAGCGTCCTGTAAC |
| *GYRA*_qRT_F          GGCGGTTTGGTTGTTTCTGG  *GYRA*_qRT_R          AACAACTCTTGCACATTTCTTGTATGG |
| *ATP-β*_qRT_F          CACCAAGAAAGGGTCTATCACC  *ATP-β*_qRT_R          GCCGTGTTGTAATGCTCCTC |
| **Gateway cloning-related (complementation and sub-cellular localization)** |
| atDjA1_N tag_AttB1 GGGGACAAGTTTGTACAAAAAAGCAGGCTGGATGTTCGGTAGAGGACC  atDjA1_N tag_AttB2  GGGGACCACTTTGTACAAGAAAGCTGGGTCTTACTGCTGGGCACATTG |
| atDjA2_N tag_AttB1  GGGGACAAGTTTGTACAAAAAAGCAGGCTGGATGTTTGGAAGAGGACC  atDjA2_N tag_AttB2  GGGGACCACTTTGTACAAGAAAGCTGGGTCTCACTGCTGGGCACATTG |
